# PlotMI: interpretation of pairwise dependencies and positional preferences learned by deep learning models from sequence data

**DOI:** 10.1101/2021.03.14.435285

**Authors:** Tuomo Hartonen, Teemu Kivioja, Jussi Taipale

## Abstract

Deep learning models have recently gained success in various tasks related to understanding information coded in biological sequences. Although offering state-of-the art predictive performance, the predictions made by deep learning models can be difficult to understand. In virtually all biological research, the understanding of how a predictive model works is as, or even more important as the raw predictive performance. Thus interpretation of deep learning models is an emerging hot topic especially in context of biological research. Here we describe PlotMI, a mutual information based model interpretation tool that can intuitively visualize positional preferences and pairwise dependencies learned by any machine learning model trained on sequence data such as DNA, RNA or amino acid sequence. PlotMI can also be used to compare dependencies present in training data to the dependencies learned by the model and to compare dependencies learned by different types of models that are trained to perform the same task. PlotMI is freely available at https://github.com/hartonen/plotMI.

## Introduction

Simultaneous rapid development of modern deep learning (DL) methods, computing hardware and high-throughput biological measurements has opened up avenues for successful application of DL to different problems in the fields of biology and medicine (1). DL models are especially well suited for analysis of large data sets of biological sequences as indicated for example by an early success story applying convolutional neural networks (CNNs) to learning binding motifs of transcription factors (TFs) (2) or more recent significant improvements in protein structure prediction using deep learning (e.g. (3)).

Biological interactions manifest themselves as dependencies in the data and learning these dependencies can often times explain why a DL model outperforms a traditional machine learning (ML) model such as linear regression. The internal representations of dependencies learned by a DL model are, however, typically non-intuitive and difficult to present in a human comprehensible format. This creates a demand for tools that can visualize dependencies learned by a DL model in an intuitive manner.

Traditionally DL has been viewed as a “black box” method giving state-of-the-art performance at the cost of losing intepretability of the model. In many biological problems the ability to explain the decisions made by an ML model is at least as important as the accuracy of the predictions. Thus several methods for interpretation of DL models trained on biological sequence data have been developed. In the following we highlight interpretation tools of DL models trained on biological sequence data as full review of DL interpretation efforts is out of the scope of this work.

One of the most straightforward methods to visualize features learned by the DL model is to directly visualize the convolutional filters of the first layer of the network (e.g. (2)). First layer filters have also been interpreted by constructing motif logos from parts of input sequences that result to high activation of the individual filters (4). Interpretation of the first layer filters is however not straightforward and the network architecture has been shown to greatly affect what kind of features the first layer filters tend to learn (5).

In silico saturation mutagenesis (ISM) (used e.g. in (2) and (4)) is an intuitive approach that measures changes in model output produced by simulated mutations. With ISM, the idea is to introduce all possible single position variants to an input sequence and score each variant with the trained DL model. The main drawback of ISM is that it is computationally expensive, as the whole model needs to be evalueted for each variant scored. Similarly to ISM, the so called feature attribution methods weigh individual positions in selected input samples based on internal DL model metrics such as gradients or activation levels (e.g. (6–8)), but with use of back-propagation making the computation much faster. However, methods such as DeepLIFT (7) compute the feature attribution scores against certain reference sequences and the choice of the reference sequence can affect the feature attributions. Strength of both ISM and feature attribution methods in DL model interpretation is that the feature importances and predicted effects of variants can be intuitively visualized for each input sample as a sequence logo. Concentrating on individual samples can however make it difficult to recognize more complicated features and thus approaches like motif discov-ery guided by the deep learning model (9), sampling the maximum entropy distribution around sample inputs (10) and visualization of feature maps learned by the DL model (11) have been proposed.

A model-agnostic approach of interpreting DL model predictions is to feed random input to a pre-trained model and study features found from the high-scoring subset of the input (used e.g. in (12)). Here, we propose a visualization tool PlotMI, that combines this idea with a measure of pairwise dependency between two variables, mutual information (MI) (13). We use a pre-trained DL model to score sets of biological (or other) sequences, select sequences of interest based on model predictions, and compute MI between pairwise position-specific k-mer distributions. Computing MI within sequences selected based on model predictions allows discovery of dependencies learned by the model. The advantage of this approach is that it simultaneously samples a large space of high-scoring sequences and projects the pairwise dependencies learned by the model into an easily interpretable two-dimensional visualization (14) that can highlight positions of important features as well as specific distances separating features learned by the model.

Additionally, PlotMI can also be used to compare the interaction patterns learned by the model to the patterns present in the training data. Biological data can contain different types of dependencies caused by different processes and only a subset of them may be important for the task the DL model has been trained to perform. Thus comparing the dependencies learned by the model to dependencies present in the data can offer insight into which dependencies are important for the task at hand. This type of analysis can also reveal if the model has not learned some dependencies present in the training data and thus further training or a change of model architecture might be beneficial. Similarly, PlotMI can be used to compare dependencies learned by different types of models trained to perform the same task to highlight possible reasons for different predictions. Visualization of MI in input samples filtered using a DL model highlights positions and spacings of dependencies learned by the model and can be used together with for example motif discovery to identify the interacting features. Although in this work we use PlotMI to interpret ML models trained on DNA, RNA and amino acid sequence, the implementation is agnostic to the type of the model or the sequence used to train the model.

## Materials and Methods

### Computing mutual information between position-specific k-mer distributions

Assuming an input set *S* of sequences of equal length, let us denote with *P*_*i*_(*a*) the observed frequency of k-mer *a* at position *i* in the set of sequences *S* and with *P*_*ij*_(*a, b*) the observed joint frequency of k-mer *a* at position *i* and *b* at position *j*. By k-mer we mean contiguous subsequences of length *k* in any alphabet (for example A, C, G and T for DNA). The frequencies *P*_*i*_(*a*) are

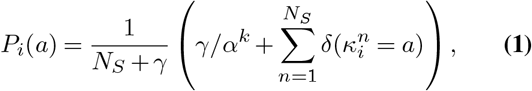

where the summation runs over the *N*_*S*_ sequences in set *S* and 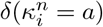 is Kronecker delta that equals to 1 only if k-mer *κ* at position *i* of sequence *n* is *a*. We add a “pseudocount mass” *γ* to the total count of k-mers accounting for the unobserved k-mers. Thus the normalization comes from the fact that there are *α*^*k*^ k-mers (where *α* is the alphabet length), and pseudocount mass *γ* is divided between all k-mers, while *N*_*s*_ k-mers are observed from the input sequences. Similarly, the observed joint frequency of k-mers *a* and *b* is computed as:

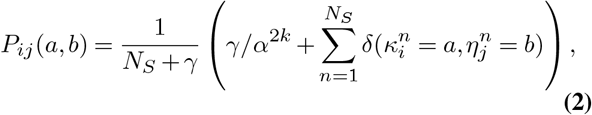

where the number of 2k-mers is *α*^2*k*^. Note that this is equivalent to counting 2k-mers with gaps. With these observed frequencies, one can compute the mutual information (originally described in (13)) between pairs of positions in the set *S* as

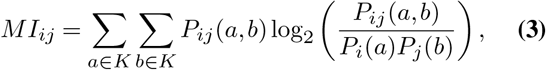

where *K* denotes the set of all k-mers and only position pairs where the k-mer distributions do not overlap with each other are considered.

Adding pseudocount to the observed k-mer counts essentially means assuming that contribution from the unobserved k-mers to the total MI is non-zero. If no pseudocount is used, the joint probability of any unobserved k-mer with any other k-mer is zero. Changing the value of pseudocount only affects the magnitude of the estimated MI values but not the overall shape of the interaction patterns highlighted by the MI plot if MI is calculated for unaligned data (see Figure S5 a-d). When visualizing models that have learned a very strict interaction, like in Figure 1 e, pseudocount drastically changes how the MI plot looks like. Figures S5 e-h show the data from Figure 1 e plotted with four different pseudocount values (10, 5, 1 & 0, respectively). Here the CACGTG-motif is present in each input sequence meaning that without adding a pseudocount it is not possible to detect any interaction involving this middle position. The dark blue “bars” corresponding to MI between the CACGTG in the middle and other positions equal exactly 0, which is less than the MI introduced by random noise between non-interacting positions. Even when using a pseudocount (Figure S5 e-g) these darker bars are still visible because the joint probability distributions of k-mers are heavily concentrated at the one observed k-mer pair at these positions. This bias is related to how the MI is estimated and vanishes when the observed k-mer counts tend towards infinity, either by increasing sample size or by increasing pseudocount.

**Fig. 1.**
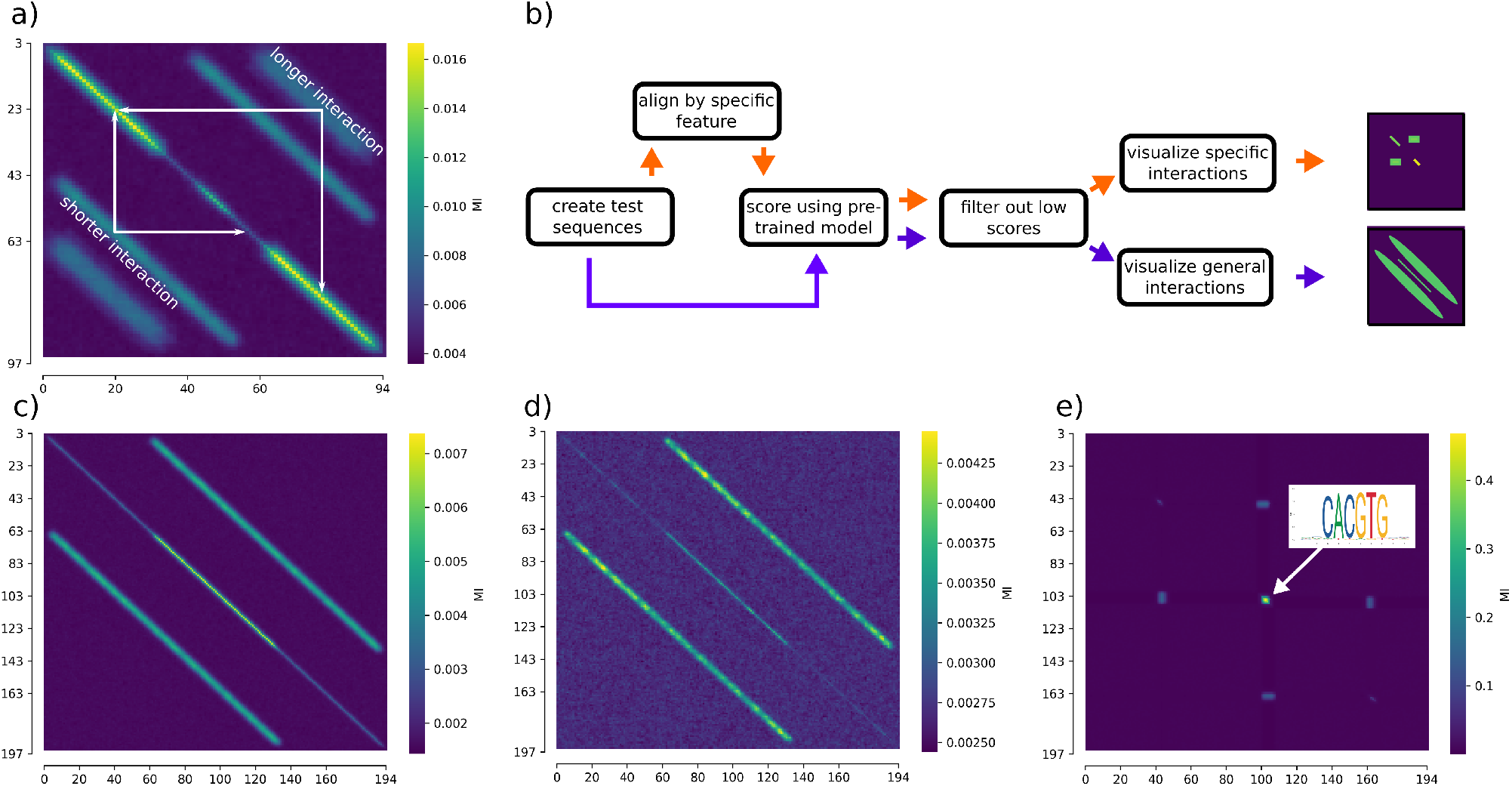
Mutual information (MI) based visualization reveals pairwise and positional dependencies. **a)** Example MI visualization of a dataset where two different pairs of transcription factor (TF) binding motifs have been embedded with different spacings (30bp and 50bp) on DNA sequences drawn from random uniform nucleotide background. MI was computed between position-specific 3-mer distributions. Signal (higher values of MI) on the main diagonal of the MI-plot highlights parts of the input sequences where adjacent positions have dependencies with each other. Signal off the main diagonal highlights pairwise longer-range dependencies. Exact positions of the interacting pairs exhibiting longer-range dependencies can be found by drawing lines parallel to the x- and y-axes towards the main diagonal, as illustrated with the white arrows here. **b)** Two possible visualization workflows using PlotMI. MI can reveal the distance between interacting features if the visualized model is trained on unaligned data and the PlotMI input sequences are also unaligned (blue arrows). If either the model is trained on aligned data, or the PlotMI input sequences have been aligned based on a feature used by the model, MI can reveal the exact positions of the interacting features (orange arrows). The values at the x and y axes indicate positions along the input sequences. Panels d and e illustrate how alignment of input data based on a feature learned by a convolutional neural network (CNN) model alters MI plot. **c)** MI-plot of the 50,000 positive class training data sequences for the CNN classifier containing the embedded MAX+ELF5 motif pair embedded at random positions but exactly 50bp from each other. The MI-plot highlights this spacing which is learned by the CNN model as shown in panel d. **d)** If the test data is not aligned, PlotMI will pick up interactions as lines parallel to the main diagonal of the MI matrix showing the preferred distance between the interacting features. MI-plot computed between position-specific 3-mer distributions shows that a CNN trained on a simulated dataset to classify between sequences with a specific MAX+ELF5 TF binding motif pair and sequences with either a MAX-, or ELF5-motif has learned the interaction embedded into the training data. **e)** If the test data is aligned based on a specific feature (or the model is trained on aligned data), interactions between features show off the main diagonal of the MI matrix. Motif discovery from the sequences visualized in panel e found a CACGTG-motif corresponding to the MAX-motif embedded into the training data of the CNN. Using the same CNN model to score sequences where this CACGTG-motif was embedded at position 100 shows that the CNN has learned a specific interaction with the CACGTG-motif and another motif that corresponds to the ELF5-motif embedded into the simulated training data (Figure S1 f).

To obtain a complete picture of pairwise dependencies learned by the machine learning model it is beneficial to visualize MI with and without pseudocount and to consider what the differences mean. For example Figure S5 h, plotted without a pseudocount, shows a dependency between the flanking positions corresponding to the ELF5 motifs mirroring the fact that in the training data there is always only one ELF5 motif embedded per sequence. This interaction is much more difficult to see with pseudocount. On the other hand considering only the MI estimated without a pseudocount will lead to missing interactions with features that are always or almost always present in the input sample.

Estimation of MI from finite samples is a non-trivial problem and different ways of estimating MI are thoroughly discussed in (15). The assumptions made here in estimating MI are 1) when the observed sample count of a k-mer approaches zero, the estimated probability of observing this k-mer approaches a non-zero constant and 2) *P*_*i*_(*a*) = ∑_*j,b*_ *P*_*i,j*_(*a, b*). In our application, the exact values of MI are not important as the main goal is to visualize the interaction patterns learned by the ML model. We have observed that setting *γ* = 5 *× N*_*s*_ usually produces good visualizations and all figures shown here have been plotted using this, unless otherwise stated. Value of pseudocount can be set by the user.

Selecting the length of k-mers to use in the MI analysis is somewhat dependent on the type of the sequence used to train the models and requires some domain knowledge on what is a typical scale for interactions in that type of data. For example, in DNA at the regulatory regions of the genome such as promoters, the interactions are typically between TF binding sites, and setting *k* = 3 can capture both interactions within TF binding motifs, that are typically around 5-15 bp long, and between parts of different binding motifs further away from each other. Figures S6 a-d show the sequences used in Figure 1 d plotted with values of *k* between 1 and 4, and Figures S6 e-h the same for the sequences used in 1 e, respectively. In this case, the same dependencies are observed regardless of k-mer length used to compute MI. However, for *k* = 4 the data starts to already get sparse (29,442 sequences in Figures S6 a-d and 13,331 sequences in Figures S6 e-h) and we see a similar effect than when not using pseudocount in Figure S5 (in Figure S6, *γ* = 5 *× N*_*s*_ for all panels). On the other hand, interactions in protein structures are typically between individual amino acid residues, making *k* = 1 a suitable choice for protein sequence. Current implementation of PlotMI has been mainly tested with k-mer lengths up to 3.

### Measuring similarity of pairwise k-mer distributions

The similarity measures of probability distributions used in Figure S1 were computed as follows. The Jensen-Shannon distance was computed using the SciPy (16) (version 1.6.3) function spatial.distance.jensenshannon, defined as:

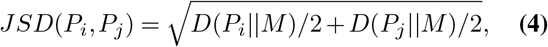

where *M* = (*P*_*i*_ + *P*_*j*_)*/*2 and *D*(*X* ||*Y*) is the Kullback-Leibler divergence (17) between distributions *X* and *Y*. Bhattacharyya distance (18) was computed as:

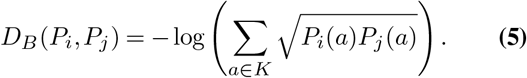

Hellinger distance (19) was computed as:

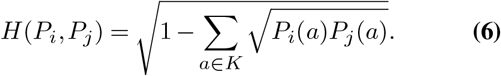

### Visualization of dependencies in unaligned versus aligned sequences

PlotMI can visualize dependencies regardless of whether the input sequences are aligned based on some feature or not. As demonstrated in Figure 1, the same dependencies will look different in aligned versus unaligned sequences. In aligned sequences, a specific spacing between feature A used to align the sequences and any feature B will result into a dependency where both the positions of A and B are fixed. If the sequences are not aligned, the exact same interaction will result into a dependency, where only the spacing between the features is fixed, but the relative position of the feature pair within the sequences varies. If the model has not been trained on aligned sequences, it can still be used to score sequences aligned based on a feature learned by the model and dependencies within the aligned sequences can then be vsiualized using PlotMI as demonstarated in Figure 1.

### Data and models

For the examples based on simulated data shown in Figure 1, the data was generated as follows: random DNA sequences were generated using randomReads script (https://github.com/hartonen/randomReads) by sampling each base separately from random uniform background distribution. Motifs were embedded to the sequences using the option to sample the embedded sequences from the corresponding position frequency matrices (PFMs). This means that the motif-sequence embedded to each background sequence was drawn at random from the corresponding PFM. The PFMs used for MAX, ELF5, KLF12 and HOXA9 transcription factors were generated using the multinomial method and have been previously published (20). The PFM matrices are given in Supplementary Table 1. Order of the motifs within embedded motif pair was drawn at random, as was the position of the motif pair within the sequence. The spacing between the drawn motif-sequences (distance from end of motif-sequence 1 to start of motif-sequence 2) in the MAX-ELF5 pairs was 30 bp (see Figure S1 a). The spacing of the drawn motif-sequences in the KLF12-HOXA9 pairs was 50 bp (Figure S1 b). Logomaker (21) Python package was used to visualize the PFMs in Figures S1a-b.

Convolutional neural network (CNN) classifiers were trained for binary classification task where class 1 sequences contain random uniform 200bp DNA sequences with a pair of motif sequences drawn at random from the MAX and ELF5 PFMs inserted at random positions exactly 50bp apart from each other (50 bp spacing meaning the distance between the end of the first motif-sequence and the start of the second motif-sequence), in random order. Class 0 sequences are also 200bp long random uniform DNA where a single motif sequence drawn at random from either MAX or ELF5 motif is inserted at random position. This data was divided at random into training (100,000 sequences), validation (20,000 sequences) and test (20,000 sequences) sets where classes are balanced. The CNN models consist of convolutional modules with a 1D convolutional layer followed by batch normalization, ReLu activation and a dropout layer. The following hyperparameters of the model were optimized using grid search: number of convolutional modules: {1, 2, 3, 4, 5, 6, 7}, number of convolutional filters per module: {32, 64}, dropout rate: {0.3, 0.4, 0.5}. Batch size 128 and convolutional kernel size 5 were used. The convolutional modules used dilated convolution (22) with dilation rate of the ith layer *i*^2^. The final layer is a dense layer of two nodes with sigmoid activation. The models were implemented with Keras (https://keras.io/) using TensorFlow 1.14.0 backend (23) and trained using Adam optimizer with default parameter values. Training was stopped once binary accuracy on validation data did not improve within 200 epochs. Final model was selected based on best binary accuracy on validation data (0.990) and had the following hyperparameters: number of convolutional modules = 7, number of filters per module = 64. dropout rate = 0.3. The final model achieved area under precision-recall curve (AUprc) score 0.997 on unseen test data.

Motif discovery from 3,037 sequences obtaining a class 1 probability >0.9 (a subsample of sequences shown in Figure 1 b) was conducted using the MEME-ChIP (24) tool by setting motif width between 6 and 25, looking for maximum of 4 motifs and otherwise using default parameters. The best motif reported by the STREME-tool (25) from MEME-ChIP was compared to database motifs with TOMTOM (26), and the best matching motif (JASPAR (27) ID: MA0825.1, shown in Figure 1 e) was selected for generating the data for Figure 1 e. Full MEME-ChIP results can be found from Supplementary File 1. Sequence logo of the sequences shown in the MI-plot in Figure 1 e was made using WebLogo (28).

Next, we trained similar CNN models to classify between human genomic promoters and non-promoter sequences (Figures 2 a-c). The 29,598 human transcription start site (TSS) positions and the *±*50bp sequences around them for hg38 reference genome were downloaded from the Eukaryotic Promoter Database (EPD) (29). These sequences were divided at random into training (20,000 sequences), test (4,799) and validation (4,799) datasets and used as class 1 for the binary classifiers. Balanced class 0 sets were created by randomly shuffling the class 1 sequences while preserving their 3-mer frequencies using the fasta-shuffle-letters tool from MEME Suite (30). The CNN architecture and training was similar to what is described above. Hyperparameters were optimzed using grid search over the following values: number of con-volutional modules: {1, 2, 3, 4, 5, 6}, number of convolutional filters per module: {64, 128}, dropout rate: {0.3, 0.4, 0.5}. Final model was selected based on best binary accuracy on validation data (0.882) and had the following hyperparameters: number of convolutional modules = 6, number of fil-ters per module = 128. dropout rate = 0.4. The final model achieved AUprc score 0.949 on unseen test data.

**Fig. 2.**
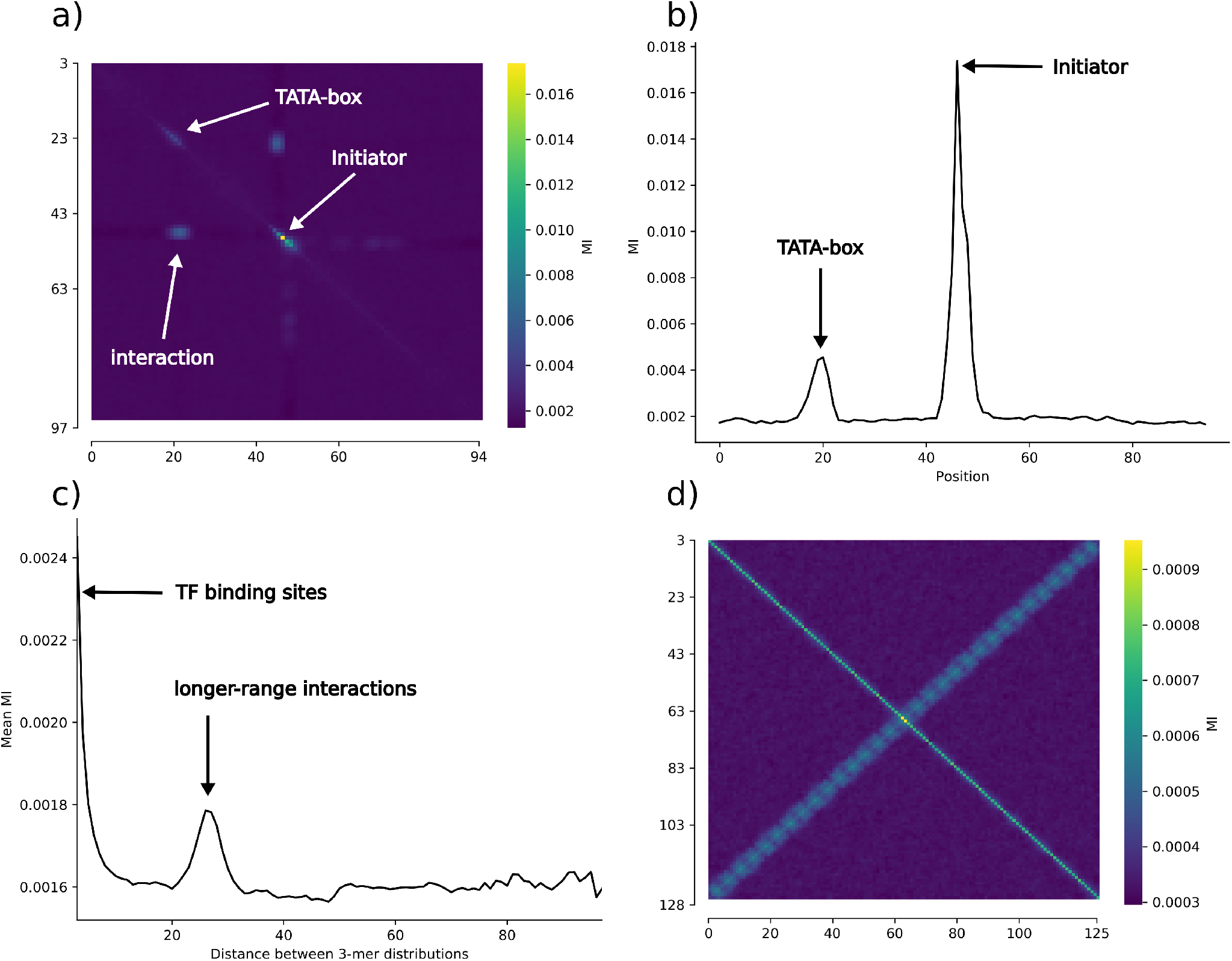
PlotMI-visualizations of machine learning models trained on genomic DNA sequences. **a)** PlotMI reveals that a CNN classifier trained on 100 bp long sequences centered at human genomic transcription start site (TSS) positions has learned an interaction between the TSS and TATA-box region around 30 bp upstream from TSS. **b)** Mutual information (MI) at the main diagonal of the MI-matrix shows exact positions of the highest local MI features. **c)** Average MI of each diagonal of the MI-matrix further illustrates that the CNN model of human promoters has learned both the short-range dependencies corresponding to neighboring 3-mer distributions (TF binding motifs) and longer range interactions corresponding to the dependencies between individual features. **d)** PlotMI visualization of N-score model trained to distinguish 131 bp long nucleosome binding DNA from non-nucleosomal DNA (31) from yeast genome shows that N-score has learned a periodic interaction pattern where 3-mer distributions at positions separated from each other by multiplicatives of a fixed period are dependent on each other (See also Figure S3). The learned interaction is symmetric relative to the middle position of the 131 bp sequences such that the strongest pairwise dependencies are observed between positions same distance away from the middle position, but to opposite directions. Both MI-analyses were done using position-specific 3-mer distributions.

The final CNN models used to filter the data shown in Figures 1 d-e and Figures 2 a-c are deposited to Zenodo along with their respective training, validation and test data sets under DOI: 10.5281/zenodo.5508698.

As an additional example of models trained on genomic DNA, we studied dependencies learned by a non deep learning model trained to recognize nucleosome favoring sequences, N-score (31) (Figure 2 d). We downloaded the pre-trained N-score implementation from https://bcb.dfci.harvard.edu/~gcyuan/nscore.zip and used it to score random synthetic DNA sequences as is without modifications.

In Figure 3, we demonstrated the use of PlotMI with models trained to classify RNA sequences to those that bind a specific RNA-binding protein (RBP) and to those that do not. We downloaded the DeepBind (2) (version 0.11) models from http://tools.genes.toronto.edu/deepbind/download.html and used them to score random synthetic RNA sequences as is without modifications. The 10 million synthetic RNA sequences were generated from random uniform background distribution using the randomReads script. The DeepBind IDs of the used models are: D00123.001 (MSI1) and D00198.001 (RBMS1).

**Fig. 3.**
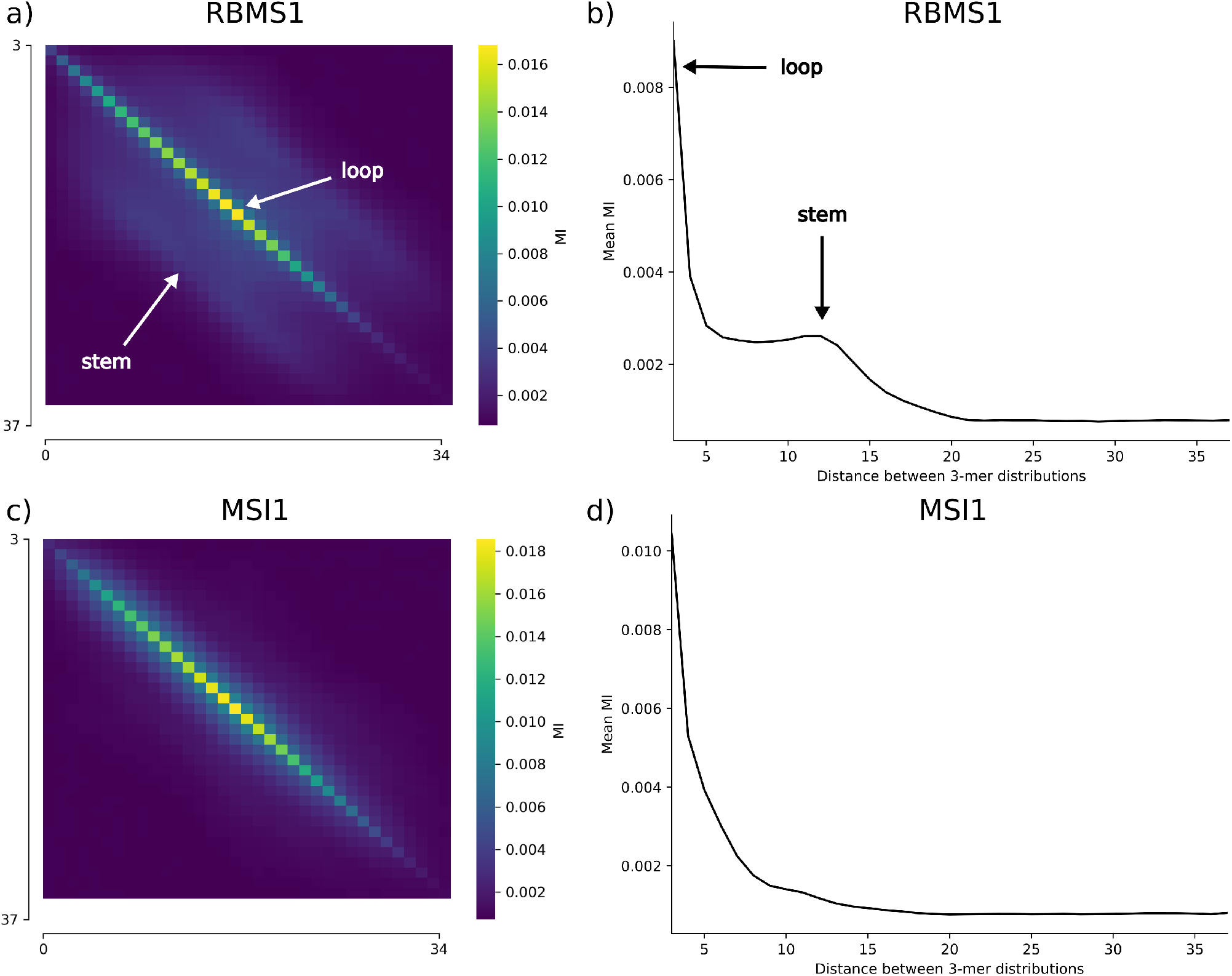
PlotMI-visualizations of selected DeepBind (2) RNA-binding protein (RBP) models. PlotMI analysis shows that some of the DeepBind RBP models have learned longer-range dependencies, consistent with the RBP recognizing a stem-loop structure, while others only consider very short-range dependencies when predicting which RNA sequences the RBP binds to. **a & b)** MI-plot and mean MI of each diagonal of the MI-plot, respectively, for the RBMS1 DeepBind model, shows MI also off the main diagonal indicating that this model has learned longer-range dependencies. **c & d)** MI-plot and mean MI of each diagonal of the MI-plot, respectively, for the MSI1 DeepBind model, shows MI signal only at the main diagonal corresponding to the model using short-range dependencies only.

As a final example, we used PlotMI to interpret dependencies learned by models that were trained to predict the fitness of amino acid sequences to function as functional proteins (Figure 4). The pre-trained protein fitness models (32) were downloaded from the GitHub repository https://github.com/gitter-lab/nn4dms and used to score random synthetic amino acid sequences as is without modifications. The 10 million synthetic amino acid sequences were generated from random uniform background distribution using the randomReads script. The contact map for GB1 protein domain (Protein Data Bank ID: 2QMT (33)) was computed with a custom BioPython (34) script using distances between *α*-carbons of each amino acid residue. The wild type GB1 sequence was the same used in (32): “MQYKLILNGKTLKGETT-TEAVDAATAEKVFKQYANDNGVDGEWTYDDATK-TFTVTE”. The fitness scores from the different models for this sequence were: -0.07492757 for graph convolutional neural network (GCN), -0.21727669 for sequence convolutional neural network (CNN) and -2.4499352 for linear regression (LR).

**Fig. 4.**
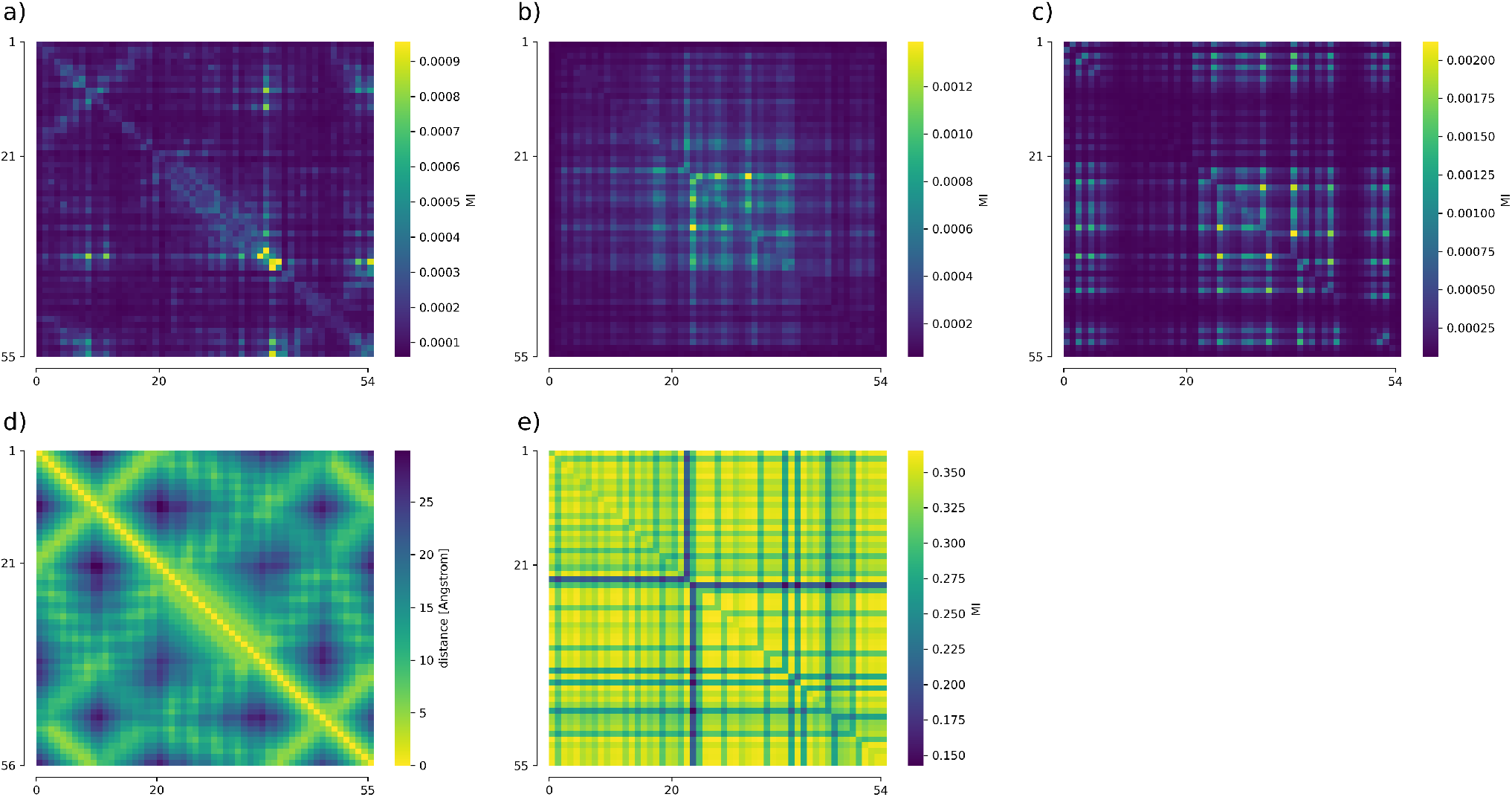
PlotMI-visualizations of models predicting protein fitness reveal different types of dependencies learned by different models. MI-plots created using position-specific 1-mer distributions showing the pairwise dependencies learned by **a)** the GCN model, **b)** the CNN model and **c)** the linear regression model from (32). **d)** Contact map of GB1 protein domain derived from 2QMT PDB structure (33) that is used in training of the GCN model. Color indicates distance between each pair of alpha carbons. Note that darker colors mean longer distance so that the heatmap can be readily compared with the MI-plots, where darker colors mean lower pairwise MI. **e)** MI-plot showing the top 26,804 (5%) training set amino acid sequences according to fitness in the deep mutational scanning experiment of the GB1 protein domain (35). Notice that only the GCN model (panel a), that has been trained with both the mutational scanning (panel e) and the structural (panel d) data, has learned bulk of the interactions important for the structure of the functional protein.

## Results

Mutual information (MI) measures dependence between two probability distributions. When MI is computed between k-mer distributions at all pairs of positions in a set of sequences (e.g. DNA or amino acid), it will highlight those position pairs where certain pair, or pairs, of k-mers appear at different frequencies than expected assuming no interactions (see for example (14)). Figure 1 a) shows an example of an MI-plot where pairwise MI was calculated between all position-specific 3-mer distributions in a synthetic example dataset containing two spiked-in dependencies between DNA sequence motifs. The shorter range interaction was created by spiking in a pair of transcription factor (TF) binding motifs, MAX and ELF5, into half of the sequences at random positions with a spacing of 30 bp between the motifs (see Methods and Figure S1 a). The longer range interaction was spiked-in into the other half of the sequences, consisting of a sequence motif pair KLF12 and HOXA9 separated by a 50 bp spacing (see Methods and Figure S1b. Main diagonal of the MI-plot shows dependencies between k-mer distributions at adjacent positions, highlighting for example TF binding motifs in case of DNA sequences. Signal off the main diagonal corresponds to interactions between more distant positions. The white arrows in Figure 1 a highlight how the exact interacting positions are read from the MI-plot by drawing arrows from the off-diagonal signal to the main diagonal. Figure 1 a illustrates how the MI-plot clearly highlights the two distinct interactions present in the data, whereas metrics designed for computing the distance between two probability distributions, such as Jensen-Shannon distance, Hellinger distance or Bhattacharyya distance, do not (Figures S1 c-e).

Workflow for the analysis of pairwise dependencies learned by a deep learning (DL) model using PlotMI is shown in Figure 1 b. Details of calculation of MI are described in Methods. To illustrate the capability of PlotMI to capture interactions learned by a DL model, we created a binary classification task where class 1 consist of DNA sequences sampled at random from uniform nucleotide background distribution and where binding motifs of TFs MAX and ELF5 were embedded into random positions in each of the sequences with exactly 50 bp spacing between the motifs. Class 0 sequences have similar background, but a single instance of either MAX-, or ELF5-motif was embedded into each sequence at random position (see Methods). Figure 1 c shows an MI-plot computed between positional 3-mer distributions in the training set sequences with the embedded interaction. A convolutional neural network (CNN) was trained to classify between these two classes and subsequently used to score 10 million DNA sequences drawn from random uniform nucleotide background. All sequences obtaining a class 1 probability *>* 0.9 (29,442 sequences) were visualized with PlotMI (Figure 1 d). PlotMI visualization highlights that the CNN model has learned the embedded interaction (see also Figure S2). Strongest MI is found between positions separated by 60 bp, as this is the distance between the most defined bases in the embedded motifs.

To demonstrate how MI plot can be used to discover specific pairwise interactions, we ran STREME (25) motif discovery on the high-scoring sequences and discovered a CACGTG-motif corresponding to the MAX-motif, one of the motifs in the embedded motif pair (see Methods). We then created another set of 1 million random DNA sequences and embedded the CACGTG-motif in the middle of these sequences. The sequences were similarly scored using the CNN and sequences obtaining class 1 probability *>* 0.9 (13,331 sequences) were visualized with PlotMI. Figure 1 e shows that the CNN scores highly those sequences where another motif is approximately 60 bp off from the centered CACGTG-motif and that there is strong MI between these positions. A sequence logo of these 13,331 sequences revealed that this other motif corresponds to the ELF5-motif (Figure S1 f).

To study interactions learned from real datasets we downloaded 29,598 experimentally validated human transcription start site (TSS) positions from the Eukaryotic Promoter Database (29) and trained a CNN model to classify between *±*50 bp sequences around the TSS and shuffled versions of these sequences. We generated 1 million 100 bp long random DNA sequences with uniform nucleotide frequencies and scored them using the CNN yielding 49,261 sequences with promoter probability *>* 0.9. PlotMI visualization shows that the highest MI is found between adjacent positions right around the TSS, and that also a highly localized pairwise interaction between the TSS and the canonical TATA-box position at around 30 bp upstream from the TSS (Figure 2 a) is learned. Plotting only the MI at the main diagonal helps to pinpoint the exact positions of high MI features learned by the model (Figure 2 b). Mean MI of each diagonal of the MI-plot shows the strength of MI between 3-mer distributions as function of distance separating the distributions (Figure 2 c) illustrating that most of the pairwise dependencies learned by the CNN model are very short-ranged, with the exception of a peak corresponding to dependencies between features separated by 25-30 bp (TSS and the TATA-box).

To demonstrate the use of PlotMI with non-DL models, we used PlotMI to analyse dependencies learned by a previously published model N-score, a wavelet-based logistic regression classifier developed to predict the nucleosome binding affinity of 131 bp long sequences in yeast (31). As PlotMI visualization is not dependent on the architecture of the machine learning model, it can be used to visualize pairwise dependencies of many different types of models. We generated 2.5 million synthetic DNA sequences with uniform nucleotide background, scored them with N-score and selected the highest-scoring 10% of the sequences for visualization. Figure 2 b) shows that, interestingly, N-score has learned a dependency pattern where strongest MI outside the main diagonal is observed between positions equidistant from the middle position of the 131 bp sequences. Moreover, the strength of these dependencies varies with a period of 9 bp (Figure S3 a). Main diagonal of the MI matrix shows that all positions share MI with their adjacent positions. Interestingly, PlotMI visualization of the bottom 10% lowest-scoring sequences shows an even more prominent symmetric and periodic dependency pattern (Figure S3 b). These examples demonstrate that PlotMI can visualize different types of dependency patterns learned by different kinds of models without needing parameter or workflow adjustments.

Next, we used PlotMI to analyse dependencies learned by DeepBind deep learning models (2) trained to recognize binding sites of specific RNA-binding proteins (RBPs). In addition to recognizing linear binding motifs similar to binding sites of TFs on DNA, some RBPs are known to recognize RNA secondary structure features (see for example (36)) which can introduce longer range dependencies a model needs to learn in order to optimally recognize binding sites of such RBPs. In Figure 3 we visualize the dependencies learned by two DeepBind RBP models: RBMS1 that has been previously reported to recognize a stem-loop structure (36), and MSI1 that binds to a linear motif (36). The PlotMI analysis was conducted by generating 10 million random synthetic RNA sequences and scoring each of them with both DeepBind models. Then separately for both models, top 1% of the highest-scoring sequences were used for computing MI between positional 3-mer distributions. Figures 3 a-b show that the RBMS1 model uses pairwise dependencies up to 15 bp apart in the RNA sequence for predicting the optimal RBP binding sites, consistent with the earlier reported binding motif for RBMS1 where the bases binding the stem part of the stem-loop structure are 9-18 bp apart (36). In contrast to this, the MSI1 model has only learned local dependencies up to approximately 6 bp. This analysis shows that PlotMI can reveal additional information about how models for different RBPs make predictions that cannot be directly inferred by only examining the motif logos learned by the models.

PlotMI can also be used to highlight different dependencies learned by different models trained to do the same task. This can help understand why some more complex models are able to outperform simpler models. As an example, we downloaded linear regression (LR), sequence convolutional neural network (CNN) and graph convolutional neural network (GCN) models trained to predict insulin binding affinity (referred to as fitness in the following) of 56 amino acid long GB1 protein sequences (32). The MI-plot shown in Figure 4e shows the top 5% training set amino acid sequences according to measured fitness from the deep mutational scanning experiment in (35) used to train these models. Notably, only single and double mutants of the wild type GB1 sequence were included in the training data set, meaning that only a very small fraction of the possible sequence space, and only very near the wild type sequence, was available for the models during training. Additionally, structural graph of GB1 protein domain was used in training of the GCN model (32). Contact map showing the physical distances between each amino acid residue pair computed from this structure (Methods) is shown in Figure 4 d.

To test what dependencies each of the models have learned, we created 10 million random synthetic amino acid sequences and scored them with the three models. We choose top 100,000 (1%) of the random sequences according to the predicted fitness for visualization with PlotMI (Figures S4 d-f show the MI-plots for sequences with predicted fitness higher than the predicted fitness of the wild type sequence). MI was computed between position-specific 1-mer distributions. Figures 4 and S4 d-f show that, unsurprisingly, the GCN model where the structural information was utilized during training had learned more of the important pairwise dependencies highlighted by the contact map than the other models. Interestingly, in (32) the authors report no significant difference in performance of the GCN and CNN models in predicting fitnesses of protein variants tested in deep mutational scanning experiments (Pearson correlation ≈ 0.98), even though the MI-plots show that the CNN utilizes only a small fraction of the pairwise dependencies present in the contact map. Based on this, it seems that the structural graph used in training of the GCN helps to assign realistic fitness values for proteins that are further away from the wild type sequence than any examples in the deep mutational scanning-based training data. The CNN seems to have hard time in recognizing variants with very poor fitness as it gives almost any sequence at least the same score as for the wild type (Figure S4 b) but can still predict the fitness of variants close to the wild type as well as the GCN. Also the LR model predicts that more than half of the random amino acid sequences have a higher fitness than the wild type GB1 sequence (Figure S4 c), whereas the GCN model predicts that only a few percent of the random synthetic amino acid sequences have higher fitness than the wild type (Figure S4 a). In contrast to the GCN and CNN model architectures, the LR model can only learn additive effects between positions. Still the pairwise positional dependency pattern learned by the LR model is relatively similar to the CNN. Based on the analysis with PlotMI, it seems that the limited sequence space available for these protein models during training prohibits the CNN and LR models from learning a realistic representation for the functional GB1 protein, even though the models can still make relatively good predictions for mutational effects near the wild type sequence. The PlotMI analysis shows that even though the performance of the GCN and the CNN models is similar in predicting mutational effects near the wild type sequence, the structural graph used to constrain the training of the GCN model has allowed it to learn a more generalizable model that better captures the dependencies required for the functional GB1 protein.

## Conclusions and discussion

Here we apply the well-known measure of mutual dependency of random variables, mutual information (MI), to extract information about interactions learned by deep learning (DL) models. The aim of this approach is to project the information about pairwise and position-specific dependencies learned by a DL model into a simpler model that can be intuitively visualized. This is done by feeding random input into a pre-trained model and using the model as a filter to select a subset of input samples containing features learned by the model. The high-scoring sequences can be further mutated, or aligned based on some feature discovered using e.g. *de novo* motif mining to capture higher information content features. Some high information content features may effectively not be present in a completely random sample that is sufficiently small that it can be scored with a given machine learningn (ML) model in reasonable time. Learned features visualized using random input sequences to PlotMI can be, however, used to aid in search of more complex interactions, like we show in Figures 1 d-e, where a TF binding motif discovered by scoring random input is in turn embedded into random sequences to reveal its interaction parter. Epistatic interactions can be captured by studying features present in low-scoring samples. The overall importance of features and interactions learned by an ML model and discovered using this type of exploratory analysis can then be quantified for example with the recently introduced global importance analysis (37).

Importantly, the approach described here is not limited to MI visualization, but can be used to project information learned by a DL model to any other easily interpretable model. PlotMI is a flexible tool that can be used to interpret different types of ML models from simple linear regression to complex deep neural networks. The examples provided here demonstrate usefulness of PlotMI for classification and regression tasks and for models trained on DNA, RNA and amino acid sequences. In principle PlotMI can be used to visualize pairwise dependencies learned by any ML model trained with sequence data.

Although the idea of interpreting a DL model by feeding it random input and studying features present in high-scoring input samples is not new, a general visualization tool like PlotMI that can visualize pairwise interactions in an intuitive manner has to our knowledge not been previously described. The PlotMI visualization script is freely available and can be used in conjunction with virtually any pre-trained ML model.

We believe that the visualization approach described here will be a useful addition to the expanding toolbox of model interpretation methods available for scientists using ML to understand information encoded into DNA, protein and other types of biological sequence.

## Acknowledgements

The authors would like to thank Drs. Esko Ukkonen, Fangjie Zhu and Kimmo Palin for helpful discussions regarding the mutual information visualization, Dr. Esa Pitkänen for critical reading of the manuscript, and Dr. Leena Salmela for help with selecting a nucleosome model for visualization. The authors wish to acknowledge CSC - IT Center for Science, Finland, for computational resources.

## Author contributions

JT developed the original idea, TH implemented the PlotMI program and performed the computational experiments. TH, TK and JT planned and wrote the manuscript.

