## Supplementary Materials for "PlotMI: interpretation of pairwise dependencies and positional preferences learned by deep learning models from sequence data"

DRAFT

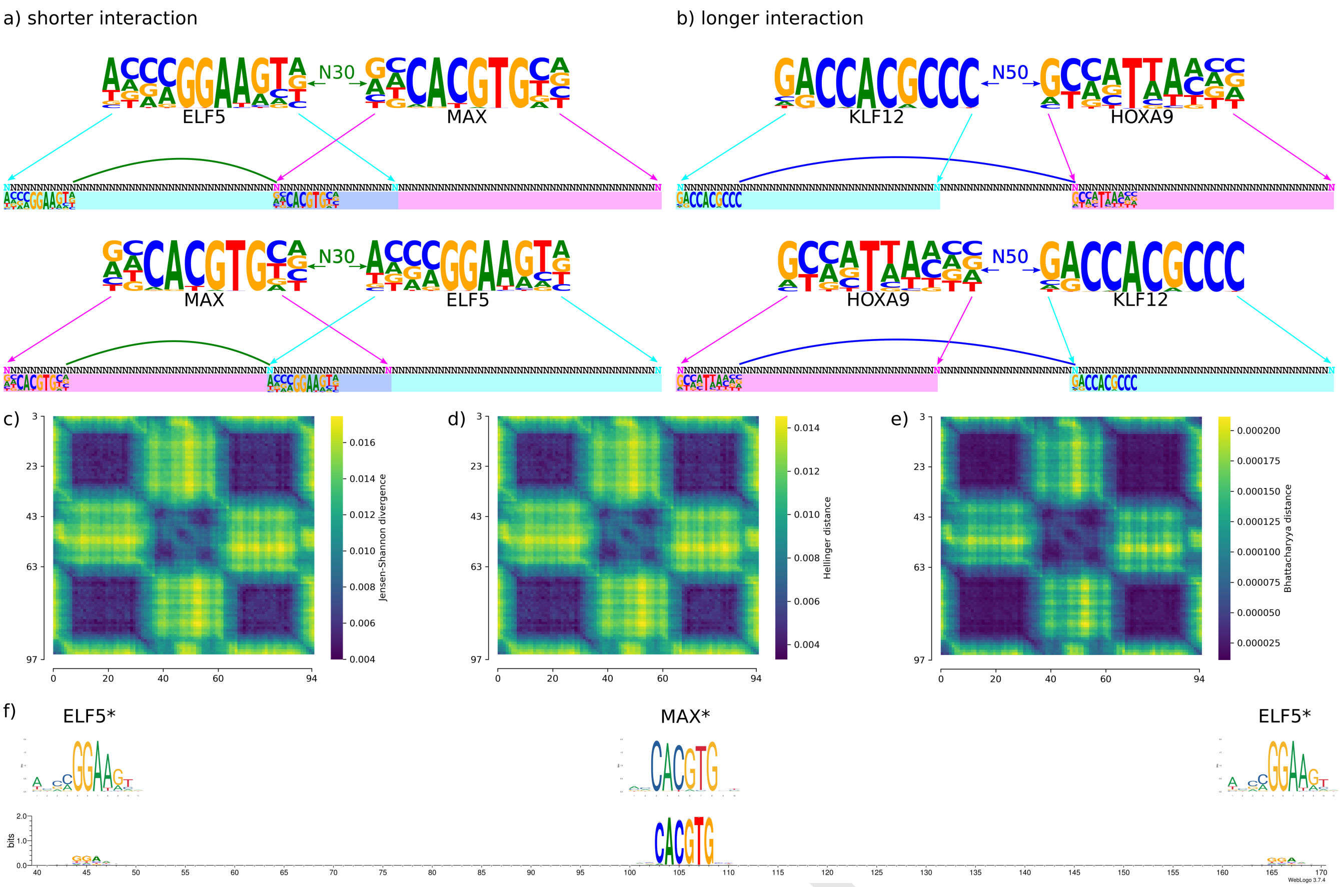


**Fig. S1. Related to Fig 1: Mutual information based visualization reveals pairwise and positional dependencies. a-b)** Schematic representation of how the example spiked-in dataset used to demonstrate MI-plot in Figure 1 a was constructed. The two interactions were spiked-in by embedding the MAX-ELF5 transcription factor (TF) motif pair (a) into half of the sequences and the KLF12-HOXA9 motif pair into the other half (b). For the MAX-ELF pair, the motifs were separated by 30 bp, meaning a 30 bp gap between the end of one motif and the start of the second. Order of the embedded motifs was decided at random as described in Methods. The KLF12 and HOXA9 motifs were separated by 50 bp. The cyan and magenta arrows and shadings indicate the possible positions for the corresponding motifs. Notice that for the shorter spacing illustrated in panel a, the possible positions of the motifs overlap in the middle of the 100 bp long sequences. Spacing between the motifs is shown with green (30 bp, panel a) and blue (50 bp, panel b). **c-e)** Visualization of a dataset where two different pairs of TFs have been embedded with different spacings (see description of the dataset from panels a-b) using Jensen-Shannon distance (c), Hellinger distance (d) and Bhattacharyya distance (e). Darker colors indicate higher similarity between positional k-mer distributions. Existence of two interactions with distinct spacings cannot be detected using measures of similarity between positional k-mer distributions. **f)** Bottom: position weight matrix (PWM) constructed from the 200bp long sequences used in MI-plot in Figure 1 e. Top: the short PWMs correspond to the motifs embedded in the training data of the CNN and are shown for comparison confirming that the CNN model has learned the interaction between the strongest bases of the embedded motifs and that the MI analysis captures this interaction. Asterisk (*) after the motif names denotes that instead of PFMs shown in panel a, position weight matrices (PWMs) are shown here, generated using flat mononucleotide background.

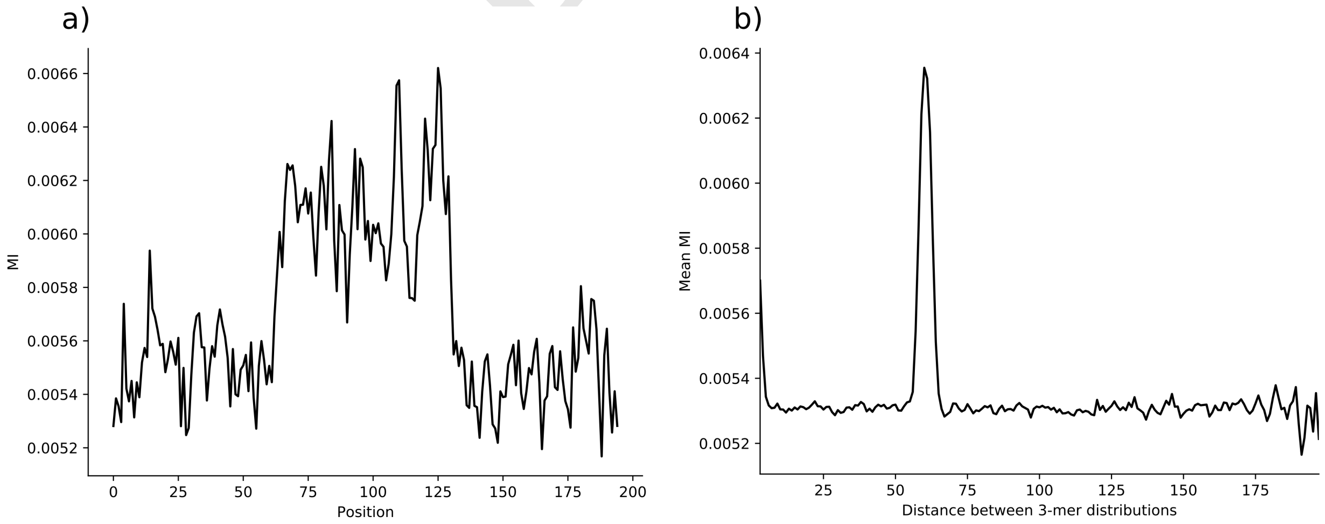


**Fig. S2. Related to Fig 1: Mutual information based visualization reveals pairwise and positional dependencies. a)** Mutual information at the main diagonal of MI-plot in Figure 1 d. **b)** Mean mutual information at each diagonal of MI-plot in Figure 1 d shows that the CNN model has learned an interaction between features that are separated by approximately 60 bps corresponding to the distance between the most defined bases in the embedded motif pair.

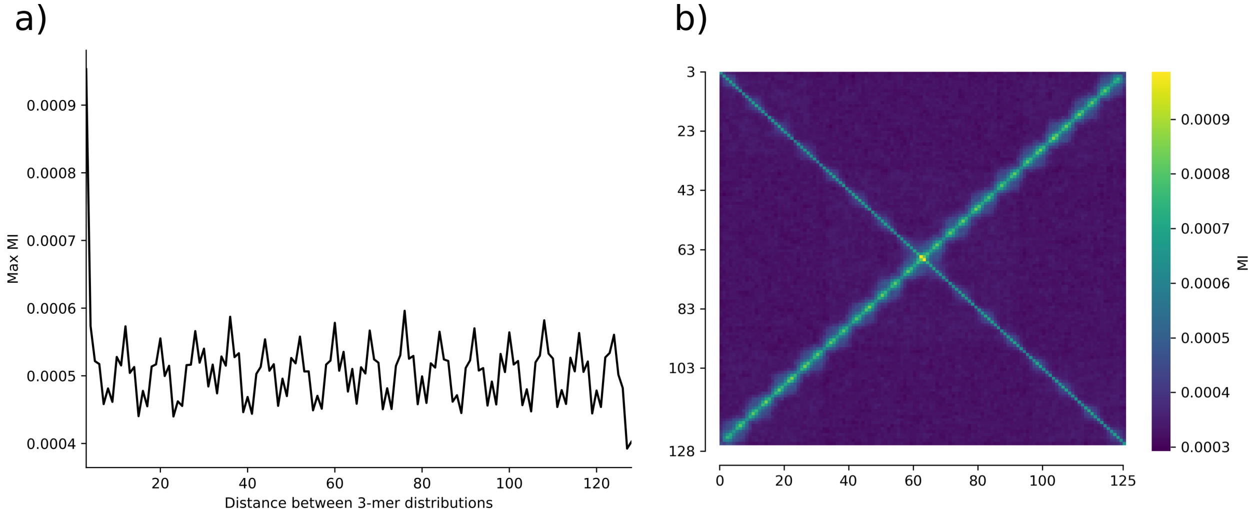


**Fig. S3. Related to Fig 2: PlotMI-visualization of machine learning models trained on genomic DNA sequences. a)** Maximum MI of each diagonal of the MI-plot describing dependencies learned by N-score (Figure 2 d) confirms that the position pairs with highest pairwise MI (outside the main diagonal) follow a 9 bp periodic pattern. **b)** PlotMI visualization of low-scoring (bottom 10 %) random 131bp long sequences scored with N-score shows that sequences not predicted to be occupied by nucleosomes show a similar periodic and symmetric dependency pattern as the high-scoring sequences shown in Figure 2 d.

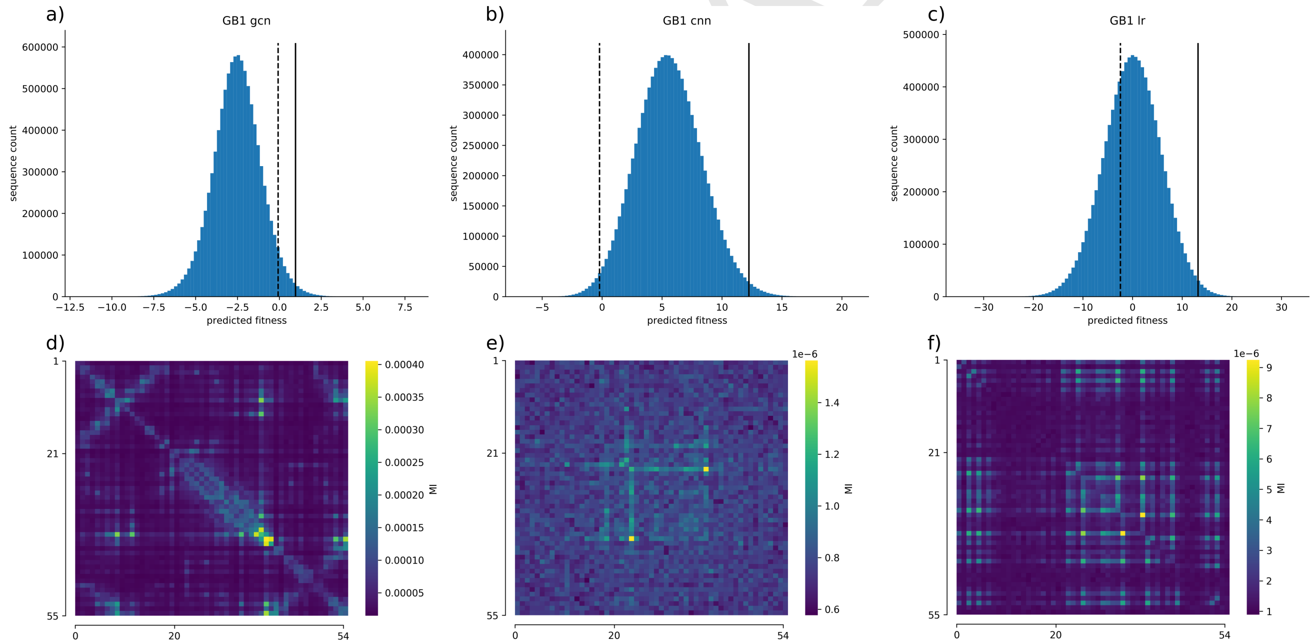


**Fig. S4. Related to Fig 4: PlotMI-visualizations of models predicting protein fitness reveal different types of dependencies learned by different models.** Histograms showing the distributions of predicted protein fitnesses of the 10 million random synthetic amino acid sequences scored by the GB1 **a)** GCN, **b)** CNN and **c)** LR models, respectively. The dashed black vertical lines indicate the fitnesses of the wild type GB1 protein domain amino acid sequence predicted by the model and the continous black vertical lines indicate the cut-off of 1% highest-scoring random amino acid sequences. MI-plots generated using the random synthetic amino acid sequences that obtain higher predicted fitness than the wild type GB1 sequence are shown in panels **d)** for GCN (447,040 sequences), **e)** for CNN (9,867,706 sequences) and **f)** for LR (6,646,479 sequences) models.

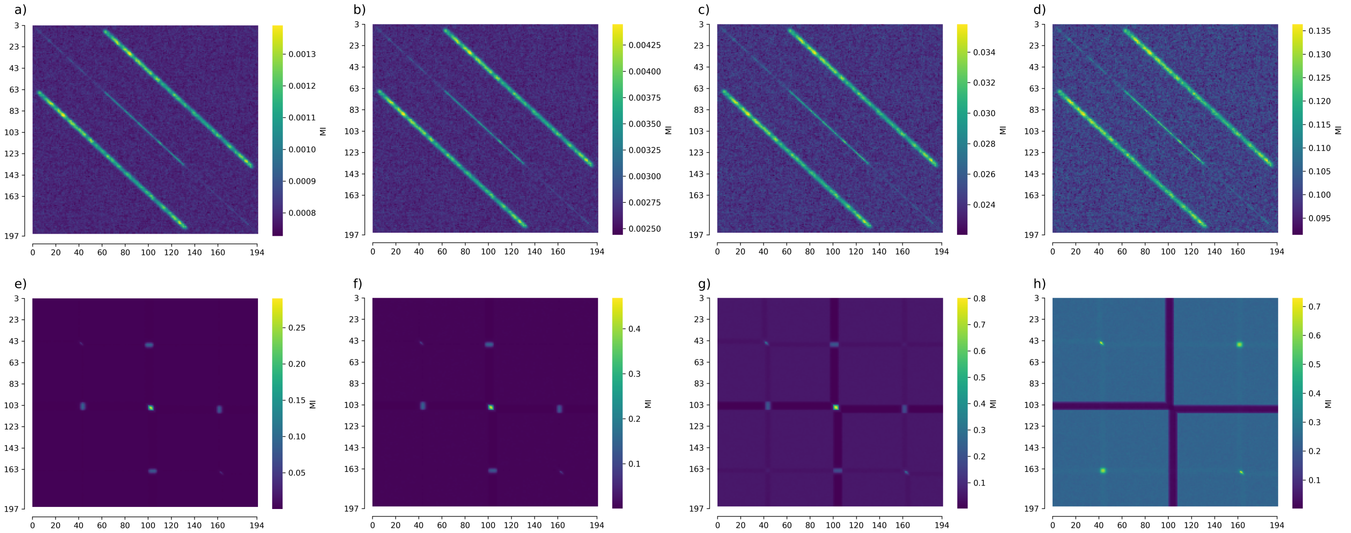

**Fig. S5. The effect of pseudocount on the MI-plot. a)** Non-aligned data, $\gamma = 10 \times N_s$. **b)** Non-aligned data, $\gamma = 5 \times N_s$ (same figure as Figure 1d). **c)** Non-aligned data, $\gamma = 1 \times N_s$. **d)** Non-aligned data, $\gamma = 0$. **e)** Aligned data, $\gamma = 10 \times N_s$. **f)** Aligned data, $\gamma = 5 \times N_s$ (same as Figure 1e). **g)** Aligned data, $\gamma = 1 \times N_s$. **h)** Aligned data, $\gamma = 0$.

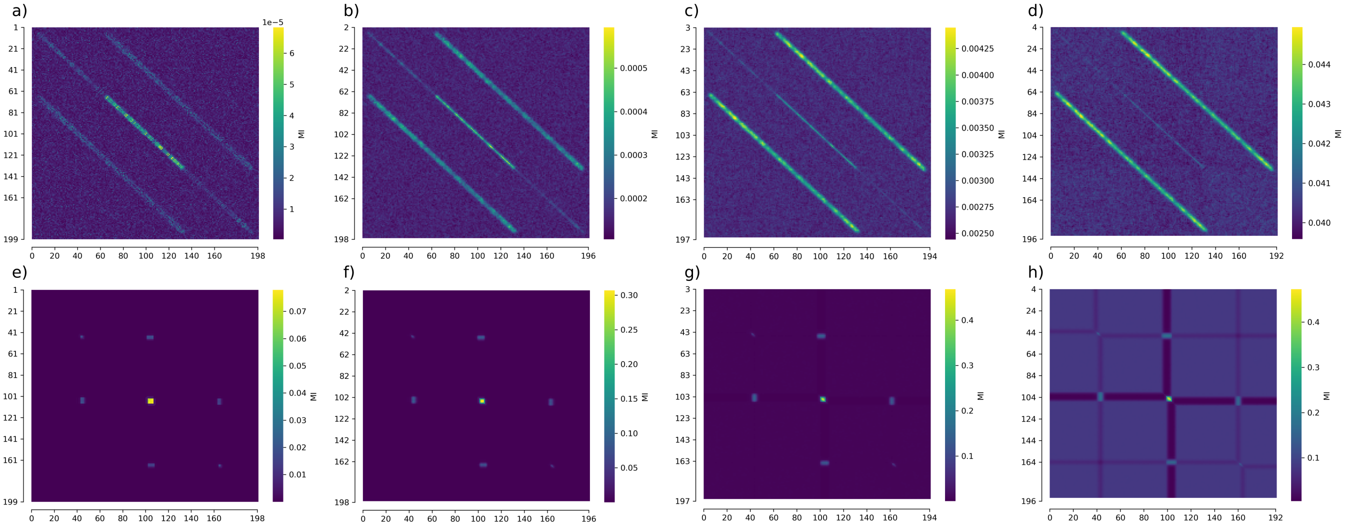

**Fig. S6. The effect of k-mer length on MI-plot. a)** Non-aligned data, k=1. **b)** Non-aligned data, k=2. **c)** Non-aligned data, k=3 (same as Figure 1d). **d)** Non-aligned data, k=4. **e)** Aligned data, k=1. **f)** Aligned data, k=2. **g)** Aligned data, k=3 (same as Figure 1e). **h)** Aligned data, k=4. Pseudocount $\gamma = 5 \times N_s$ was used in all panels.
