## Supplementary File 1 for "PlotMI: interpretation of pairwise dependencies and positional preferences learned by deep learning models from sequence data": centrimo.html

CentriMo Results

[close ]

[close ]

[
close ]

[
close ]

Each "motif probability curve" shows the (estimated) probability of the
**best** match to a given motif occurring at a given position in the
input sequences. This estimated probability is based only on sequences that
contain at least one match with  
than the  threshold defined for this motif,
and is the maximum likelihood estimate of the conditional probability shown below.

Points (X,Y) on the plot are:  
  Y = Pr(best match occurs at position X | sequence contains a match)

**Note:** The plots are smoothed according to the function
selected from the "Smoothing" menu on the right. Setting the smoothing
window size to 1 turns off smoothing.

If a negative dataset has been supplied then two curves are drawn for
each motif, one for each dataset. The distribution of the motif in the
primary dataset is plotted with a single unbroken curve, whereas the distribution in
the negative dataset is plotted with a dashed curve.

[
close ]

This shows a listing of all motifs currently plotted on the graph.

The color used to plot a motif can be changed by clicking on the
color swatch next to the motif you want to change, followed by clicking
on the color swatch you wish to swap it with.

[
close ]

These are extra colors you may use for plotting motifs.

Click on the color swatch next to one of the above motifs, then click
on one of these "unused color" swatches to change the color of the
motif's plot.

[
close ]

These options change the display of the graph.

Smoothing:
:   Allows selection of the smoothing function applied to the graph.

    The weighted moving average option uses weights shaped as an isosceles
    triangle where the central point (or points in an even sized window)
    get the maximum weight.

    The moving average simply weights all points in the smoothing window
    equally.

    **Note:** Setting the smoothing window size to 1 turns off
    smoothing.

Window
:   The window size used to smooth the graph. The larger the smoothing
    window size, the smoother the graph, at the cost of hiding detail.

    Below a smoothing window size of 10, thinner lines are used on the
    graph to allow more detail to be visible.

    **Note:** Remember to press "return" or "enter" after changing
    the number in the input box in order to see the effect of the new
    smoothing window size.

Legend
:   Choose to display/disable the on-graph legend. The legend can be
    moved by clicking on the graph.

Negative Sequences
:   Choose whether to plot the motif probability curve(s) for the
    negative sequences (if provided). The curve(s) are plotted as dashed
    lines, using the same color as the corresponding curve for the positive
    sequences.

Zoom
:   Drag a range on the graph to zoom into that section. Clicking
    "Undo Zoom" will return the view to the previously displayed part of the
    graph and clicking "Center on 0" will move the view so 0 is in the
    center.

Download EPS
:   Download the graph that you are currently viewing as an
    encapsulated postscript (EPS) image. EPS images are scalable making them
    suitable for publication.

[
close ]

List only enriched motifs that meet the selected filter criteria below.

**Selected motifs are always listed**; deselect all motifs first by clicking on
the "X" above the color swatches if you wish to filter all motifs.

To filter on "ID" or "Name", you can enter any Javascript regular
expression pattern. See here
for documentation on Javascript regular expression patterns.

[
close ]

Sorting is applied after filtering where possible (the exception being
the "Top" filter) so the filters applied will affect the sort. You can
choose the motif sorting feature using the "Motifs:" menu.

If CentriMo is searching for locally enriched regions (not just centrally
enriched regions), then multiple regions may be found per motif, and
the "Regions:" menu will also be displayed. In this case,
CentriMo first sorts all regions using the feature
shown in the "Regions:" menu, and then it sorts the highest-ranked
region of each motif according to the feature shown in the "Motifs:" menu.

Unless you check the box next to the "Regions:" menu, it will automatically
show the same feature as the "Motifs:"
menu (or "*E*-value" if a motif-only feature is chosen in the "Motifs:" menu).

**Note:**The motif *p*-value shown in the plot legend will always be for
the region with the lowest *p*-value, and therefore may not match the value
shown in the table "*p*-value" column
when the "Regions:" menu is not set to "*p*-value".

[
close ]

[
close ]

[
close ]

[
close ]

[
close ]

The expected number motifs that would have least one
region as **comparatively** enriched for
best matches to the motif as the reported region in the
**positive** sequences compared with the **negative**
sequences.

The Fisher *E*-value is the (one-sided) *p*-value of
the one-sided Fisher's exact test that **at least** as many best matches
in the region in the positive sequences that contain at least
one match, multiplied by the number of motifs in the input database(s).
The Fisher's exact test *p*-value is corrected for the number
of regions and score thresholds tested ("Multiple Tests").

Fisher's exact test assumes that the probability that the best match
(if any) falls into a given region is the same for all
positive and negative sequences.

[
close ]

[
close ]

[
close ]

The Matthew's Correlation Coefficient (MCC) gives a measure of the ability
of the motif to discriminate the positive sequences from the negative sequences:

MCC = [TP\*TN - FP\*FN] / [(TP + FP) \* (TP + FN) \* (TN + FP) \* (TN + FN)]

where

:   TP is the number of positive sequences with a best match in the reported region,
:   FP is the number of negative sequences with a best match in the reported region,
:   TN is the number of negative sequences without a best match in the reported region, and
:   FN is the number of positive sequences without a best match in the reported region.

MCC ranges from -1 to +1, where a +1 result indicates that the occurrence
of the best match to the motif in the reported region perfectly discriminates positive
sequences from negative sequences.

[
close ]

[
close ]

[
close ]

The number of **negative** sequences where the **best** match to
the motif falls in the reported region. This value is rounded but the
underlying value may contain fractional counts.

**Note:** This number may be less than the number of **negative**
have a best match in the region. The reason for this is that a sequence may
have many matches that score equally best. If *n* matches have the
best score in a sequence, 1/*n* is added to the appropriate bin
for each match.

[
close ]

The number of sequences containing a match to the motif
 the 
threshold ("Score Threshold").

[
close ]

This is the  threshold
()
for determining if a sequence contains a match to this motif.

[
close ]

The number of **negative** sequences containing a match to the motif
above the score threshold. When score optimization is enabled the
score threshold may be raised higher than the minimum.

[
close ]

The probability that any tested region in the **negative**
sequences would be as enriched for best matches to this motif according
to the Binomial test.

Use the filter to display only motifs differentially enriched in both
datasets (low *p*-value and high negative *p*-value).

[
close ]

The maximum probability that the best match occurs at any single sequence position.
If the smoothing window size ("Window:", to right of graph) is set to "1", then this is value is
the maximum value of the match-probability curve.

[
close ]

**Concentration** is defined as the total probability of all the positions in the central
region whose width is the same as the size of the "smoothing window".
You can change the size of the smoothing window using the "Window:"
input field in the **Graph** options section, above.
(A value of "NaN" indicates that the smoothing window size is too
large for the motif.)

The "concentration" of the motif sites in the central window
can somtimes be more informative than the *E*-value. For example,
in some ChIP-seq datasets, motifs for cofactors show more significant
enrichment overall (smaller *E*-value), but are less
concentrated in a small (20 to 50bp) window than the motif for
the ChIP-ed transcription factor. In such cases, you may wish
to sort the motifs by Concentration, using the **Sort**
menu, below.

[
close ]

[
close ]

Location of the center of the most enriched region.

[
close ]

The text box lists the sequence identifiers for sequences that
 **all** the
selected motifs.

The "Intersection" subheading gives the number of identifiers in the
text box and their percentage out of the total number of input sequences.

The "Union" subheading lists the number and percentage of
sequences that have at least one of their best matches in the most
significant region of **any** of the selected motifs and their
percentage out of the total number of input sequences.

Note that the number of sequences with a match to a given motif in
its best region may be larger than the value of "Region Matches". This is because
a sequence may have multiple equally best matches and in that case a
fractional match count is assigned to each of them when "Region Matches" is computed.

[
close ]

When more than one significant, non-overlapping region is found,
they can be shown (and hidden again) by clicking the arrow.

By default the regions are sorted by *E*-value, but this can be
changed by the menu on the right of the page.

[
close ]

Sequence position where the (unsmoothed) match-probability curve for this motif
attains its maximum. Set the smoothing window size ("Window:", to right of graph) to
"1" to see the unsmoothed match probability curve.

[
close ]

### CentriMo

#### Local Motif Enrichment Analysis

For further information on how to interpret these results please access
https://meme-suite.org/meme/doc/centrimo-output-format.html.
To get a copy of the MEME software please access
https://meme-suite.org.

If you use CentriMo in your research, please cite the following paper:  

Timothy L. Bailey and Philip Machanick,
"Inferring direct DNA binding from ChIP-seq",
*Nucleic Acids Research*, **40**:e128, 2012.
[full text]

Motif Probability Graph
  |  
Enriched Motifs
  |  
Input Files
  |  
Program information
  |  
Results in TSV Format  
  |  
Sequence position vs. number of matches for each motif

### Javascript is required to view these results!

### Your browser does not support canvas!

#### Results

###### Motif Probability Graph ()

###### Options

###### Plotting

###### Unused Colors

###### Graph

Smoothing:

Moving Average
Weighted Moving Average

Window:

Legend:

Disabled
Enabled (click on graph to move)

X-axis:

Position of Best Site in Sequence
Distance of Best Site from Sequence Center

Negative sequences:

Not plotted
Plotted as dashed line

Zoom:

###### Enriched motifs (*E*-value ≤ using )

| ☒ | Database | ID | Alt ID | Consensus | Concentration | *E*-value | Fisher *E*-value | *p*-value | Negative *p*-value | MCC | Region Center | Region Width | Region Matches | Sequence Matches | Negative Region Matches | Negative Sequence Matches | Max Probability | Max Probability Location | Multiple Tests | Score Threshold | Other Regions |
| --- | --- | --- | --- | --- | --- | --- | --- | --- | --- | --- | --- | --- | --- | --- | --- | --- | --- | --- | --- | --- | --- |

###### Matching sequences (out of )

###### Union: 0 sequences (0%).

###### Intersection: 0 sequences (0%).

###### Filter & Sort

###### Filters

Top

Database is

ID matches

Alt ID matches

*E*-value ≤

Fisher *E*-value≤

Region Width ≤

Negative set *E*-value ≥

###### Sort

Motifs:

Regions:

###### Columns to display

Show Database

Show ID

Show Alt ID

Show Consensus

Show Concentration

Show *E*-value

Show *p*-value

Show Fisher *E*-value

Show Matthew's Correlation Coefficient

Show Region Center

Show Region Width

Show Region Matches

Show Sequence Matches

Show Negative *p*-value

Show Negative Region Matches

Show Negative Sequence Matches

Show Max Probability

Show Max Probability Location

Show Multiple Tests

Show Score Threshold

#### Input Files

###### Alphabet

###### Sequences

| Database | Source | Sequence Count |
| --- | --- | --- |

###### Negative sequences

| Database | Source | Sequence Count |
| --- | --- | --- |

###### Motifs

| Database | Source | Motif Count |
| --- | --- | --- |

###### Other Settings

|  |
| --- |
| Objective Function |
| Convert Motifs to Different Alphabet? |
| Motif Pseudo-Counts |
| Required sequence length |
| Site Scoring Method |
| Score Threshold |
| Optimize Score Threshold? |
| Minimum Region Size |
| Maximum Region Size |
| Strand Handling |
| Plotting of Matches on Negative Strand |
| Sequence IDs Included in Output? |

###### CentriMo version

(Release date: )

###### Command line summary
