## Supplementary File 1 for "PlotMI: interpretation of pairwise dependencies and positional preferences learned by deep learning models from sequence data": fimo.html

FIMO Results


---

|  |  |  |  |  |
| --- | --- | --- | --- | --- |
| **Database and Motifs** | **High-scoring Motif Occurences** | **Debugging Information** | **Results in TSV Format** | **Results in GFF3 Format** |

  
  


---
