## Supplementary File 1 for "PlotMI: interpretation of pairwise dependencies and positional preferences learned by deep learning models from sequence data": meme-chip.html

MEME-ChIP Results


[close ]

[close ]

[
close ]

[
close ]

This is a link to the motif in the output of the particular motif
discovery (e.g., MEME) or motif enrichment (e.g., CentriMo) program that
reported it.

[
close ]

This is the significance of the motif according to the particular motif
discovery (e.g., MEME) or motif enrichment (e.g., CentriMo) program that
reported it.

Follow the link under the "Discovery/Enrichment Program" column for
more information on how the significance value was derived.

[
close ]

Motifs reported by a motif discovery program (e.g., MEME) are compared
with known motifs in a motif database specified by the user. This column
lists the (up to) three most similar motifs. Only known motifs with
TOMTOM similarity E-values of less than 1.0 to the discovered motif will
be shown here. Clicking any of these links will show the TOMTOM results
where all alignments can be viewed.

Motifs reported by a motif enrichment program (e.g., CentriMo) list the
motif's name and a link to the motif's entry on the database website if it
is available.

[
close ]

This graph shows the distribution of the best matches to the motif in
the sequences as found by a CentriMo analysis.

The vertical line in the center of the graph corresponds to the center
of the sequences.

Clicking on a motif's graph will take you to the CentriMo output with
that motif selected for graphing.

[
close ]

Clicking here will show you all the motifs found by motif discovery or
motif enrichment analysis that are significantly similar to the reported
motif.

The additional motifs are shown aligned with the reported motif,
sorted in order of significance of the motif according to the
particular motif discovery (e.g., MEME) or motif enrichment
(e.g., CentriMo) program that reported it.

To cluster the motifs MEME ChIP does the following:

1. Start with no groups and all significant reported motifs.
2. Run TOMTOM with all significant reported motifs to determine
   pairwise similarity.
3. Group Highly Similar Motifs---

   While ungrouped motifs:

   Select most significant ungrouped motif.

   This is called the "seed" motif for the group and we will call the
   E-value of its seed motif the group's "significance".

   Form a new group from the seed motif and all other motifs that
   are not yet in a group and who are strongly similar to the seed
   motif (default: TOMTOM E-value ≤ 0.05).
4. Merge Groups---

   For each group (most significant to least significant), merge it with
   any less significant group if all of its motifs are weakly similar to
   the first group's seed motif (default: TOMTOM E-value ≤ 0.1).

[
close ]

Clicking here takes you to the CentriMo motif enrichment analysis with
the results for this all the motifs in this group.

[
close ]

This lists links to related content, which may include:

- Motif Spacing Analysis--SpaMo results using this **discovered** motif as
  the "primary" motif, and each of the discovered motifs and
  motifs in any motif databases specified to MEME-ChIP as
  potential "secondary" motifs. SpaMo reports the secondary
  motifs whose occurrences are enriched at particular distances relative
  to the primary motif's occurrences in the input sequences.- Motif Sites in GFF3--FIMO results showing the positions of occurrences
    of this **discovered** motif in the input sequences in GFF3
    format. If the input sequences to MEME-ChIP have FASTA headers following
    the UCSC style ("chromosome\_name:starting\_position-ending\_position"),
    and the chromosome names are in UCSC (not ENSEMBL) format,
    the GFF3 output will be suitable for uploading to the UCSC Genome Browser
    as a custom track.

[
close ]

### MEME-ChIP

#### Motif Analysis of Large Nucleotide Datasets

For further information on how to interpret these results please access
https://meme-suite.org/meme/doc/meme-chip-output-format.html.  
To get a copy of the MEME software please access
https://meme-suite.org.

If you use MEME-ChIP in your research, please cite the following paper:

Motifs
  |  
Programs
  |  
Input Files
  |  
Program information
  |  
Summary in TSV Format 
  |  
Motifs in MEME Text Format


### Javascript is required to view these results!

### Your browser does not support canvas!


#### Motifs

**The significant motifs
(E-value ≤ )
found by the programs MEME, STREME and CentriMo;
clustered by similarity and ordered by E-value.**

Expand All Clusters
Collapse All Clusters

#### Programs

#### Input Files

###### Alphabet


###### Motifs

###### MEME-ChIP version

(Release date: )

###### Command line summary
