## Supplementary File 1 for "PlotMI: interpretation of pairwise dependencies and positional preferences learned by deep learning models from sequence data": spamo.html

SpaMo Results

[
close ]

The database of the primary motif.

[
close ]

The ID of the primary motif followed by the alternate ID in brackets if it has one.

[
close ]

The logo of the primary motif.

Sections of the motif with a gray background have been trimmed and
were not used for scanning.

[
close ]

The number of secondary motifs found that had significant spacings in
the tested region.

[
close ]

The list of secondary motifs found that had significant spacings in
the tested region.

[
close ]

The name of the sequence database.

[
close ]

The last modified date of the sequence database.

[
close ]

The number of sequences in the sequence database.

[
close ]

The number of sequences in the sequence database which were excluded
because they were shorter than twice the margin plus the primary motif
length.

[
close ]

The number of sequences in the sequence database which were excluded
because they contained large runs of ambiguous symbols (normally
wildcard masking) that could bias the results.

[
close ]

The number of sequences in the sequence database which were excluded
because no match to the primary motif could be found at a distance to
the edges larger than the margin.

[
close ]

The number of sequences in the sequence database which were excluded
because they were largely identical to other sequences when aligned on
the primary motif site.

[
close ]

The number of sequences which were scanned with the secondary
motifs.

[
close ]

The name of the motif database derived from the file name.

[
close ]

The date that the motif database was last modified.

[
close ]

The number of motifs loaded from the motif database. Some motifs may
have been excluded.

[
close ]

The number of motifs with significant *E*-values whose
significant spacings were not considered too similar to those of another motif.

[
close ]

The number of motifs that while having significant spacings were less
significant than another motif that matched
most of the same sites.

[
close ]

This checkbox ensures the row stays visible after a filter operation
that would normally hide it.

[
close ]

The ID of the secondary motif.

[
close ]

The alternate name of the secondary motif.

[
close ]

The ID of the secondary motif followed by the alternate ID in brackets if it has one.

[
close ]

The name of the cluster to which this secondary motif belongs.
SpaMo assigns each secondary motif to a cluster, and names the
cluster after the motif in it with the most significant spacing.
SpaMo assigns two secondary motifs to the same cluster if the matches
in their most significant spacings (from the primary motif) overlap substantially.
Clustering is controlled by the -joint and -overlap options.

[
close ]

The *E*-value is the lowest *p*-value of any spacing of the
secondary motif times the number of secondary motifs.
It estimates the expected number of *random* secondary motifs that
would have the observed minimum *p*-value or less.

[
close ]

The gap between the primary and secondary motifs for the most significant spacing.

[
close ]

The strand and position of the secondary motif relative to the primary motif for the most significant spacing.

[
close ]

The minimum score accepted as a match to either the primary or secondary motif. This value
can greatly affect the results of SpaMo. If it is too high, there will be no
matches to the primary motif. If too low, sequences with non-significant matches
to the primary and/or secondary motif will reduce the effectiveness of the spacing analysis.

[
close ]

The distance either side of the primary motif site which makes up the region that
can contain the secondary motif site. Additionally it is the minimum gap between
the primary motif site and the edge of the sequence. These constraints mean that
input sequences shorter than the trimmed length of the primary motif plus two
times the margin size can not be used by SpaMo.

[
close ]

A histogram showing the counts for the orientation with the best spacing.

The significant spacings are highlighted in red.

[
close ]

The primary motif is used as the reference point for all spacing
calculation.

Sections of the motif with a gray background have been trimmed and
were not used for scanning.

[
close ]

The secondary motif occurs at the spacings relative to the primary
shown in the histogram below.

Sections of the motif with a gray background have been trimmed and
were not used for scanning.

[
close ]

The regions matching the secondary motif in the sequences with
the given spacing are used to construct a motif. The logo
for this "inferred" motif is shown aligned with that of the actual
secondary motif.

The inferred secondary motif logo should closely resemble that
of the secondary motif. If it does not, this may suggest that
the observed spacing may actually be due to the enrichment of a motif
that differs from the secondary motif.

You can download the inferred secondary motif by moving the mouse
cursor over the logo and clicking "Download as MEME motif". You can then
use this downloaded motif as an input to Tomtom to see what other known
motifs it may resemble.

[
close ]

These are the sequence logos created by aligning all of the
sequences with the significant motif spacing. Alignments
are centered on the match to the primary motif and done
separately for each of the quadrants that contribute to the
significant spacing. The logos extend in both
directions (up to) 10 positions past the maximum region
considered in the significance tests.

**Note 1:** If you don't
see the complete logo(s), you can use the scroll bar underneath
the Alignment window. If you don't see a scroll bar and are
on a Mac, you can turn on scroll bars by clicking on the Apple Icon
at the top left of your terminal and clicking:
`System Preferences/General/Show scroll bars/Always`.

**Note 2:**These logos are useful for detecting cases where highly
similar regions (such as DNA repeats) are present
among the sequences with the significant motif spacing.
Such cases may indicate that the spacing is due
to recent duplication events rather than to
a functional biological relationship between
the primary and secondary motifs. Ideally, the
regions around the primary and secondary motifs should
have low information content and their logos in the
alignment should closely match their motifs.

[
close ]

This table shows the details of the significant spacings between the primary motif
and the secondary motif currently selected in the "Secondaries" section, below.
Click on a row in this table to select a particular spacing for detailed analysis.

Gap
:   is the space between the primary and secondary motifs where a value
    of zero means there is no space between them. Note that if a motif
    has had low information content areas trimmed off this is the
    gap to the first untrimmed position.

Orientation
:   is the combination of quadrants used. Possible values are:
    individual quadrants (up+, up-, dn+, dn-) which are important
    when neither motif is palindromic; the diagonally combined quadrants
    (up+/dn-, up-/dn+) which are important when only the primary motif is
    palindromic; the vertically combined quadrants (up+/up-, dn+/dn-) which
    are important when only the secondary motif is palindromic; and all
    quadrants combined together (all) which is important when both motifs are
    palindromes.

*P*-value
:   is the probability of the observed number (or more) sequences
    having the observed spacing between the primary and secondary motif,
    adjusted for multiple tests. The number of multiple tests is the
    number of spacing bins (the number of bars in one quadrant of the histogram)
    times the number of combinations of quadrants (nine)
    tested for significance.

[
close ]

The histogram below shows the frequency of spacings from the primary
motif to the secondary motif.

The two quadrants on the left show spacings where the secondary motif
is upstream of the primary motif and the two quadrants on the right show
spacings where the secondary motif is downstream of the primary motif.

The two quadrants on the top show spacings where the secondary motif
is on the same strand as the primary motif and the two quadrants on the
bottom show spacings where the secondary motif is on the opposite strand
to the primary motif.

Histogram bars highlighted pink are part of one of the listed significant
spacings. This feature can be disabled by unchecking the "highlight all"
option under the spacings.

Histogram bars highlighted red are part of the currently selected
significant spacing. This feature can be disabled by unchecking the
"highlight selected" option under the spacings.

[
close ]

The selected orientation graph shows the combined quadrants from the
selected spacing with a zoomed view that only shows the portion of the
graph for which significance testing was performed.

Histogram bars highlighted pink are one of the listed significant
spacings for this orientation. This feature can be disabled by unchecking
the "highlight all" option under the spacings.

The histogram bar highlighted red is the currently selected
significant spacing. This feature can be disabled by unchecking the
"highlight selected" option under the spacings.

[
close ]

This causes a file named `spamo_contr_seqs.txt` or
`spamo_contr_seqs.bed`
to be downloaded. The file contains the **contributing sequence IDs**
for each significant spacing.

Each group of sequence IDs begins
with a comment line containing (1) the rank of the spacing, (2) the name of the file
that would contain the sequence IDs if you had used the
"Contributing Sequence IDs Download" function for a single spacing,
and (3) the *p*-value of the spacing. (**Note:** See the help bubble for
"Contributing Sequence IDs", below, for the format and meaning of the file names.)

The sequence identifiers will be as they appear in the input sequence file
(Plain) or in UCSC Genome Browser format (BED),
depending on which file you choose to download.

[
close ]

This lists the sequence identifiers of the subset of sequences
that contain the significant motif spacing. You can choose either
the original sequence ID format (Plain) or UCSC Genome Browser format
(BED) using the menu below.

These identifiers can be cut-and-pasted into other
programs for further analysis (e.g., Genome Ontology
analysis or location analysis in the case ChIP-seq peak regions).

You can also download the identifiers using the "Download" link below.
They will be placed in a file with name:

`seqs_<prim>_with_<scnd>_g<gap>_o<orient>`

and extension `.txt` if you choose "Plain Format"
or with the extension `.bed` if you choose BED format.
The fields in brackets in the file name have the following meanings:

| name | meaning |
| --- | --- |
| `<prim>` | the ID of the primary motif |
| `<scnd>` | the ID of the secondary motif |
| `<gap>` | the width of the spacing |
| `<orient>` | an integer code denoting the orientation of the spacing. |

The orientation codes are:

| orientation code | enriched quadrant(s) |
| --- | --- |
| 0 | up+ |
| 1 | dn+ |
| 2 | up- |
| 3 | dn- |
| 4 | up+/up- |
| 5 | up+/dn- |
| 6 | up-/dn+ |
| 7 | dn+/dn- |
| 8 | all |

[
close ]

Click on a row in this table to select one of the significant
secondary motifs for detailed analysis.
The details of the significant spacings between the primary motif
and the secondary motif you select here will be displayed in the
table and plots above.

[
close ]

Specify which secondary motifs to display in the Secondaries table
by checking one or more of the tick boxes below and then entering
filter criteria. Then click "Update" to refresh the view of the
Secondaries table.

[
close ]

Specify the order in which secondary motifs are displayed in the Secondaries table
by selecting a sorting criteria in the menu below.
Then click "Update" to refresh the view of the
Secondaries table.

[
close ]

### SpaMo

#### Spaced Motif Analysis Tool

For further information on how to interpret these results please access
https://meme-suite.org/meme/doc/spamo-output-format.html.  
To get a copy of the MEME software please access
https://meme-suite.org.

If you use SpaMo in your research please cite the following paper:

Primary Motifs  |  Sequence Database  |  Secondary Motif Databases  |  Spacing Analysis  |  Inputs and Settings  |  Program information  |  Results in TSV Format

  |  Contributing Sequence IDs [Download Plain] [Download BED]

#### Primary Motifs

Next Top

| Database | Name | Preview | Significant Secondaries | List |
| --- | --- | --- | --- | --- |

###### Alphabet

#### Sequence Database

Next Previous Top

| Name | Last Modified | Contained | Too Short | Too Masked | No Primary | Too Similar | Used |
| --- | --- | --- | --- | --- | --- | --- | --- |

#### Secondary Motif Databases

Next Previous Top

| Name | Last Modified | Number of Motifs | Motifs Significant | Motifs Redundant |
| --- | --- | --- | --- | --- |

#### Settings

Next Previous Top

|  |  |
| --- | --- |
| Match Score Threshold |  |
| Margin size |  |
| Width of histogram bins |  |
| Significance computed up to this distance |  |
| Secondary match handling | Count only the best secondary match above the score threshold Count all secondary matches above the match score threshold |
| Maximum allowed sequence identity |  |
| Odds ratio for redundancy heuristic |  |
| Bin *p*-value cutoff |  |
| Secondary motif *E*-value cutoff |  |
| Overlapping bases for redundancy check |  |
| Fraction of sites for redundancy check |  |
| Pseudocount added to motifs |  |
| Bit threshold for trimming motif edges |  |
| Primary and secondary motif alphabets | Converting secondary alphabet to primary alphabet Primary and secondary alphabets must match |
| Random number seed |  |
| Show Advanced Settings Hide Advanced Settings | |

#### Spacing Analysis for

Next Previous Top

|  |  |  |  |  |  |  |  |  |
| --- | --- | --- | --- | --- | --- | --- | --- | --- |
| Secondary Motif: |  | Cluster: |  | E-value: |  | Best Gap: |  | Best Orientation: |

##### Primary Motif Logo

##### Secondary Motif Logo

##### Inferred Secondary Motif Logo

Download as EPS

Download as MEME motif

Download as EPS

|  | Alignment Logo |
| --- | --- |
| Up+ | Download as EPS Download as MEME motif |
| Up- | Download as EPS Download as MEME motif |
| Dn+ | Download as EPS Download as MEME motif |
| Dn- | Download as EPS Download as MEME motif |

Download as EPS

  

##### Spacings

| Gap  Gap | Orientation Orientation | *p*-value *p*-value |
| --- | --- | --- |

Highlight:
Selected

All

##### Overview Graph

Download as EPS

##### Selected Orientation Graph

Download as EPS

##### Contributing Sequence IDs ()

  

Plain Format
BED Format
Download

##### Secondaries

###### Filter

Top

ID matches

Name matches

Cluster matches

*E*-value ≤

Gaps (ranges allowed)

###### Sort

Sort by

ID
Name
Cluster
E-value
Gap
Orientation
Spacing

| Lock | ID | Name | Cluster | E-value | Best Gap | Best Orientation | Spacings |
| --- | --- | --- | --- | --- | --- | --- | --- |

Previous Top

###### SpaMo version

 (Release date: )

###### Command line

asdf  
  
Result calculation took  seconds
