## Supplementary File 1 for "PlotMI: interpretation of pairwise dependencies and positional preferences learned by deep learning models from sequence data": streme.html

STREME Results

Help poup.

[close ]

STREME results in plain text format.

[
close ]

STREME results in XML format.

[
close ]

The name of the motif uses the IUPAC codes for nucleotides or proteins.
Letters representing multiple nucleotides are used in nucleotide motif
positions where several nucleotides are favored.

The name of the motif is <index>-<consensus>,
where <index> is the order in which the motif was found,
and <consensus> is an approximation of the motif
by an IUPAC sequence.

Read more about the MEME Suite's use of the IUPAC alphabets.

[close ]

The logo of the motif.
The rules for construction logos are given in
the Description section of the documentation for the MEME Suite utility
ceqlogo.

[close ]

The logo of the reverse complement motif.
The rules for construction logos are given in
the Description section of the documentation for the MEME Suite utility
ceqlogo.

[close ]

Show detailed information about the motif.

[close ]

Submit your motif to another MEME Suite program (see list below), or Download your motif
as a probability matrix, count matrix or MEME formatted motif, or download
a Sequence Logo for your motif.

###### Supported Programs

Tomtom
:   Tomtom is a tool for searching for similar known motifs.

MAST
:   MAST is a tool for searching biological sequence databases for
    sequences that contain one or more of a group of known motifs.

FIMO
:   FIMO is a tool for searching biological sequence databases for
    sequences that contain one or more known motifs.

GOMo
:   GOMo is a tool for identifying possible roles (Gene Ontology
    terms) for DNA binding motifs.

SpaMo
:   SpaMo is a tool for inferring possible transcription factor
    complexes by finding motifs with enriched spacings.

[close ]

The number of positive sequences matching the motif.

[close ]

The number of training set positive sequences matching the motif / the number of training set positive sequences.

Note these counts are made after erasing sites that match previously
found motifs.

[close ]

The number of training set positive sequences matching the motif.

Note these counts are made after erasing sites that match previously
found motifs.

[close ]

The number of training set negative sequences matching the motif / the number of training set negative sequences.

Note these counts are made after erasing sites that match previously
found motifs.

[close ]

The number test set positive sequences matching the motif / the number of test set positive sequences.

Note these counts are made after erasing sites that match previously
found motifs.

[close ]

The number of test set positive sequences matching the motif.

Note these counts are made after erasing sites that match previously
found motifs.

[close ]

The number of test set negative sequences matching the motif / the number of test set negative sequences.

Note these counts are made after erasing sites that match previously
found motifs.

[close ]

The mean distance from the center of the best match to the sequence center,
averaged over all training set sequences with a match.

[close ]

The mean distance from the center of the best match to the sequence center,
averaged over all test set sequences with a match.

[close ]

The Score is the unadjusted *p*-value of the motif based on the appropriate test
applied to the training set sequences.
Since the Score is not adjusted for multiple tests, it **cannot**
be used to determine the statistical significance of the motif.
The Score is used by STREME to select the best motif at each iteration.

For determining if a motif is statistically significant, you should use the
value in the P-value column. If there is no P-value column, that
means that the positive hold-out set would have been too small (fewer than 5 sequences).
For very small sequence sets, it is not practical for STREME to compute an accurate *p*-value.
In that case, you can determine if your motif is significant by running STREME twenty or more
times on shuffled versions of your positive dataset,
and seeing if the Score is always **larger** than the Score using the original sequences.
You can make shuffled sequence datasets using the MEME Suite command-line utility
fasta-shuffle-letters) if you
have installed the MEME Suite on your own computer.

The statistical test used in computing the Score is either the Fisher Exact Test,
the Binomial Test, or the Cumulative Bates distribution. (See Inputs and Settings
for the particular test being used.) The Fisher Exact Test and the Binomial Test both
measure the enrichment of the motif in the positive test sequences compared to the
the negative test sequences.
(The Binomial Test is used when the positive and negative sequences have different average lengths.)
The Cumulative Bates distribution measures the tendency
of motif to be near the center of the sequences.

[close ]

The *p*-value of the motif based on applying the appropriate statistical test
to the test set sequences. The *p*-value is an **accurate estimate** of
the statistical significance of the motif as long as the length
distributions of the positive and negative sequences are the same.

The statistical test used in computing the *p*-value is either the Fisher Exact Test,
the Binomial Test, or the Cumulative Bates distribution. (See Inputs and Settings
at the bottom of this document for the particular test being used.)
The Fisher Exact Test and the Binomial Test both
measure the enrichment of the motif in the positive test sequences compared to the
the negative test sequences.
(The Binomial Test is used when the positive and negative sequences have different average lengths.)
The Cumulative Bates distribution measures the tendency
of motif to be near the center of the sequences.

[close ]

The score threshold for determining if a potential site is a match
to the motif. The same threshold is applied when determining matches in
the training and test sequences. The threshold is in bits.

The match score of a position in a sequence is determined by converting
the motif to a base-2 log-odds matrix using the formula log2(prob[a][i]/background[a]).
Here, prob[a][i] is the probability of the letter 'a' at position 'i' of the motif,
and background[a] is the probability of the letter 'a' according to the background.

[close ]

The name of the file containing the (positive) sequences in which STREME will search for
enriched motifs.

[close ]

The name of the file containing the negative (e.g., control)
sequences relative to which STREME will look for motifs enriched
in the positive sequences, or the words "n-Order Shuffled Sequences" if
no negative sequence file was given and the negative sequences are
shuffled copies of the positive sequences.  
  
0-order shuffling preserves 1-mer frequencies (i.e., the letter frequencies),
1-order shuffling preserves 2-mer frequencies, etc.

[close ]

The name of the alphabet of the sequences.

[close ]

The number of sequences.

[close ]

The name of the alphabet symbol.

[close ]

The frequency of the alphabet symbol in the positive sequences.

[close ]

###### Details

| Train Positives | Train Positives | Train Negatives | Train DTC | Score | Test Positives | Test Positives | Test Negatives | Test DTC | P-value | Match Threshold |
| --- | --- | --- | --- | --- | --- | --- | --- | --- | --- | --- |
| /  () | /  () |  |  |  | /  () | /  () |  |  |  |  |

x

#### Submit or Download

⇧⬆

⇩⬇

Submit MotifDownload MotifDownload Logo

###### Submit to program

|  |  |  |
| --- | --- | --- |
|  | Tomtom | Find similar motifs in published libraries or a library you supply. |
|  | FIMO | Find motif occurrences in sequence data. |
|  | MAST | Rank sequences by affinity to groups of motifs. |
|  | GOMo | Identify possible roles (Gene Ontology terms) for motifs. |
|  | SpaMo | Find other motifs that are enriched at specific close spacings which might imply the existence of a complex. |

Format:

Count Matrix
Probability Matrix
Minimal MEME

|  |  |
| --- | --- |
| Format: | PNG (for web) EPS (for publication) |
| Orientation: | Normal Reverse Complement |
| Small Sample Correction: | Off On |
| Width: | cm |
| Height: | cm |

### STREME

#### Sensitive, Thorough, Rapid, Enriched Motif Elicitation

For further information on how to interpret these results please access
https://meme-suite.org/meme/doc/streme.html.

If you use STREME in your research please cite the following paper:

Discovered Motifs
  |  
Inputs & Settings
  |  
Program Information
  |  
Results in Text Format 
  |  
Results in XML Format

### Javascript is required to view these results!

### Your browser does not support canvas!

#### Description

#### Discovered Motifs

Next Top

No motifs were discovered!

#### Inputs & Settings

Previous Next Top

###### Positive Sequences

| Source | Alphabet | Sequence Count |
| --- | --- | --- |

###### Negative Sequences

| Source | Sequence Count |
| --- | --- |

###### Background

###### Other Settings

|  |  |
| --- | --- |
| Strand Handling | This alphabet only has one strand. Only the given strand is processed. Both the given and reverse complement strands are processed. |
| Objective Function |  |
| Statistical Test |  |
| Motif Selection Criterion |  |
| Minimum Motif Width |  |
| Maximum Motif Width |  |
| Background Model |  |
| Sequence Shuffling |  |
| Test Set |  |
| Word Evaluation |  |
| Seed Refinement |  |
| Refinement Iterations |  |
| Minimum Score |  |
| Refinement Match Subsets |  |
| Minimum Palindrome Ratio |  |
| Maximum Palindrome Edit Distance |  |
| Print Candidate Motifs? |  |
| Random Number Seed |  |
| Total Length |  |
| Maximum Motif P-value |  |
| Maximum Motifs to Find |  |
| Maximum Run Time |  |

Previous Top

###### STREME version

(Release date: )

###### Command line
