## Supplementary figures and images for "PlotMI: interpretation of pairwise dependencies and positional preferences learned by deep learning models from sequence data"

### logo1.png

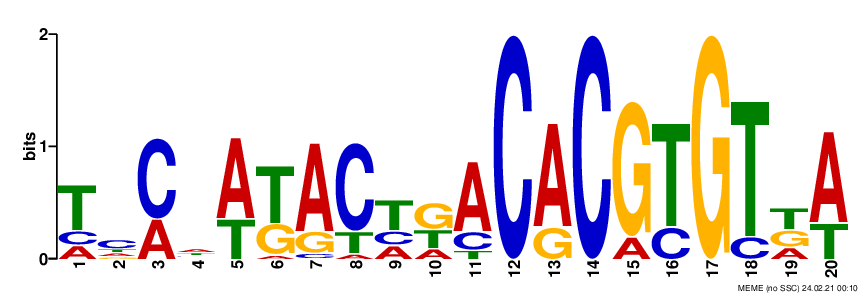

### logo2.png

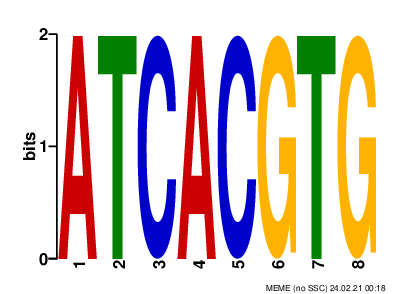

### logo3.png

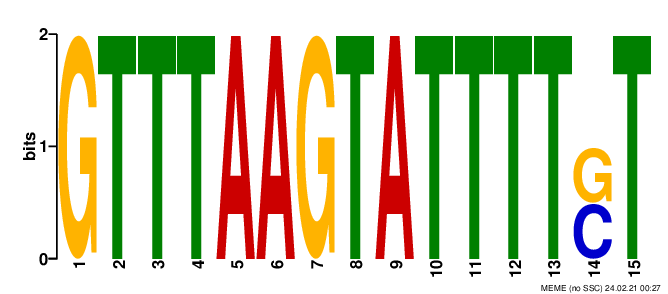

### logo4.png

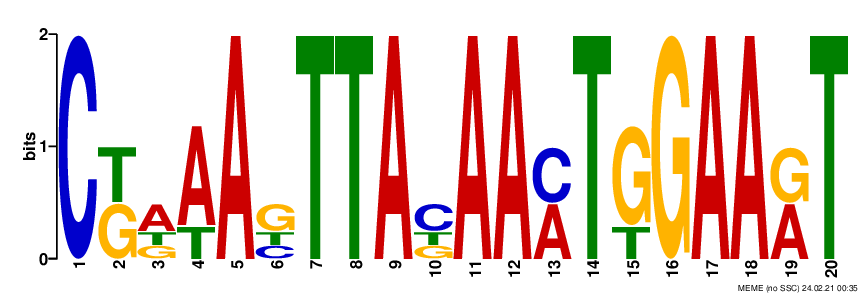

### logo_rc1.png

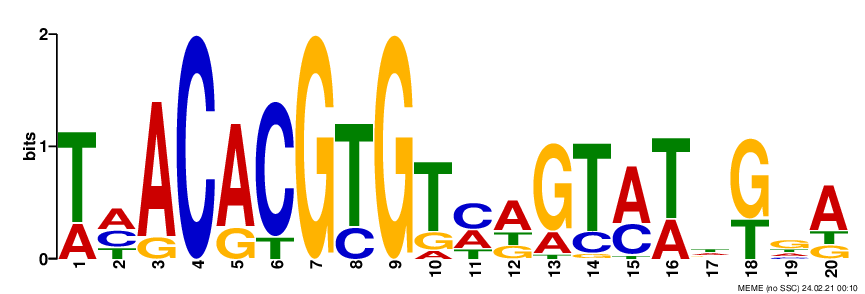

### logo_rc2.png

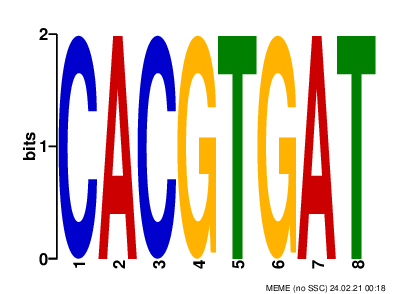

### logo_rc3.png

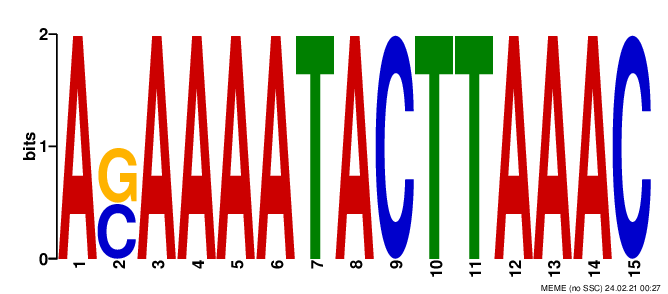

### logo_rc4.png

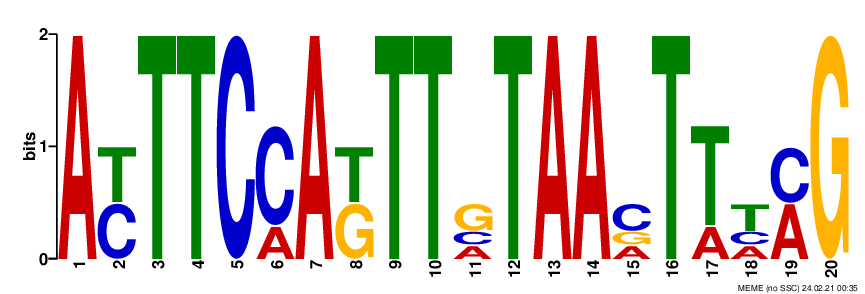
